# Carbon monoxide, a retrograde messenger generated in post-synaptic mushroom body neurons evokes non-canonical dopamine release

**DOI:** 10.1101/382127

**Authors:** Kohei Ueno, Johannes Morstein, Kyoko Ofusa, Shintaro Naganos, Ema Suzuki-Sawano, Saika Minegishi, Samir P. Rezgui, Hiroaki Kitagishi, Brian W. Michel, Christopher J. Chang, Junjiro Horiuchi, Minoru Saitoe

**Affiliations:** Tokyo Metropolitan Institute of Medical Science, 2-1-6 Kamikitazawa, Setagaya-ku, Tokyo, 1568506, Japan; Departments of Chemistry and Howard Hughes Medical Institute, University of California, Berkeley, CA, 94720, USA; Department of Chemistry, New York University, NY 10012, USA; Department of Molecular Chemistry and Biochemistry, Faculty of Science and Engineering, Doshisha University, Kyotanabe, Kyoto, 6100321, Japan; Department of Chemistry and Biochemistry, University of Denver, CO, 80208, USA

**Keywords:** Drosophila, dopamine, retgrade messenger, carbon monoxide

## Abstract

Dopaminergic neurons innervate extensive areas of the brain and release dopamine (DA) onto a wide range of target neurons. However, DA release is also precisely regulated, and in *Drosophila,* DA is released specifically onto mushroom body (MB) neurons, which have been coincidentally activated by cholinergic and glutamatergic inputs. The mechanism for this precise release has been unclear. Here we found that coincidentally activated MB neurons generate carbon monoxide (CO) which functions as a retrograde signal evoking local DA release from presynaptic terminals. CO production depends on activity of heme oxygenase in post-synaptic MB neurons, and CO-evoked DA release requires Ca^2+^ efflux through ryanodine receptors in DA terminals. CO is only produced in MB areas receiving coincident activation, and removal of CO using scavengers blocks DA release. We propose that DA neurons utilize two distinct modes of transmission to produce global and local DA signaling.

**SIGNIFICANCE STATEMENT:** Dopamine (DA) is needed for various higher brain functions including memory formation. However, DA neurons form extensive synaptic connections, while memory formation requires highly specific and localized DA release. Here we identify a mechanism through which DA release from presynaptic terminals is controlled by postsynaptic activity. Postsynaptic neurons activated by cholinergic and glutamatergic inputs generate carbon monoxide, which acts as a retrograde messenger inducing presynaptic DA release. Released DA is required for memory-associated plasticity. Our work identifies a novel mechanism that restricts DA release to the specific postsynaptic sites that require DA during memory formation.

## INTRODUCTION

Dopamine (DA) is required for various brain functions including the regulation of global brain states such as arousal and moods(Huang and Kandel, 1995; Molina-Luna et al., 2009; Yagishita et al., 2014). To perform these functions, individual DA neurons innervate extensive areas of the brain and release DA onto a wide range of target neurons through a processes known as volume transmission(Agnati et al., 1995; Rice and Cragg, 2008; Matsuda et al., 2009). However, this extensive innervation is not suitable for precise, localized release of DA, and it has been unclear how widely innervating dopaminergic neurons can also direct DA-release onto specific target neurons.

In *Drosophila*, olfactory associative memories are formed and stored in the mushroom bodies (MBs) where Kenyon cells, MB intrinsic neurons which are activated by different odors, form synaptic connections with various MB output neurons (MBONs) which regulate approach and avoidance behaviors(Gerber et al., 2004; Aso et al., 2014). Dopaminergic neurons modulate plasticity of Kenyon cell MBON synapses(Claridge-Chang et al., 2009; Aso et al., 2010; Aso et al., 2012; Liu et al., 2012). However, while there are approximately 2000 to 2500 Kenyon cells that form thousands of synapses with MBONs, plasticity at these synapses is regulated by relatively few DA neurons(Mao and Davis, 2009). This indicates that canonical action potential-dependent release cannot fully explain DA release and plasticity. We recently determined that in *Drosophila,* synaptic vesicular (SV) exocytosis from DA terminals is restricted to mushroom body (MB) neurons that have been activated by coincident inputs from odor-activated cholinergic pathways, and glutamatergic pathways, which convey somatosensory (pain) information(Ueno et al., 2017). Odor information is transmitted to the MB by projection neurons (PNs) from the antennal lobe (AL)(Marin et al., 2002; Wong et al., 2002), while somatosensory information is transmitted to the brain via ascending fibers of the ventral nerve cord (AFV). AL stimulation evokes Ca^2+^ responses in the MB by activating nicotinic acetylcholine receptors (nAChRs), and AFV stimulation evokes Ca^2+^ responses in the MBs by activating NMDA receptors (NRs) in the MBs(Ueno et al., 2013). Significantly, when the AL and AFV are stimulated simultaneously (AL + AFV) or the AL and NRs are stimulated simultaneously (AL + NMDA), plasticity occurs such that subsequent AL stimulations causes increased Ca^2+^ responses in the α3/α’3 compartments of the vertical MB lobes(Wang et al., 2008; Ueno et al., 2013). This plasticity is known as long-term enhancement (LTE) of MB responses and requires activation of D1 type DA receptors (D1Rs) in the MBs. Furthermore, while activation of D1Rs alone is sufficient to produce LTE, neither AL nor AFV stimulation alone is able to cause SV exocytosis from presynaptic DA terminals projecting onto the α3/α’3 compartments of the vertical MB lobes. Instead, exocytosis from DA terminals occurs only when postsynaptic Kenyon cells are activated by coincident AL + AFV or AL + NMDA stimulation. Strikingly, while MBs are bilateral structures and DA neurons project terminals onto both sides of MBs(Mao and Davis, 2009), SV exocytosis occurs specifically in MB areas that have been coincidently activated. Based on these results, we proposed that coincident inputs specify the location where DA is released, while DA induces plastic changes needed to encode associations. However, it has been unclear how activated Kenyon cells induce SV exocytosis from presynaptic DA terminals. Locally restricted SV exocytosis upon coincident activation of Kenyon cells requires activity of the *rutabaga* type of adenylyl cyclase, which is proposed to be a coincident detector in the MBs. Thus Kenyon cells may sense coincident activation and send a retrograde “demand” signal to presynaptic terminals to evoke local DA release(Ueno et al., 2017).

In this study, we used a *Drosophila* dissected brain system to examine synaptic plasticity and DA release, and found that coincidentally activated post-synaptic Kenyon cells generate the retrograde messenger, carbon monoxide (CO). CO is generated by heme oxygenase (HO) in post-synaptic MB neurons, and induces DA release from pre-synaptic terminals by evoking Ca^2+^ release from internal stores via ryanodine receptors (RyRs). Thus, while individual DA neurons extensively innervate the MBs, on-demand SV exocytosis allows DA neurons to induce plasticity in specific target neurons.

## MATERIALS AND METHODS

### Fly Stock maintenance

All fly stocks were raised on standard cornmeal medium at 25 ± 2°C and 60 ± 10% humidity under a 12/12 h light–dark cycle. Flies were used for experiments 1-3 d after eclosion.

### Transgenic and mutant lines

All transgenic and mutant lines used in this study are listed in supplemental Table S1. *UAS-G-CaMP3* (BDSC_32234, Bloomington Stock Center, Indiana), *LexAop-G-CaMP2* (Ueno et al., 2013) and *LexAop-R-GECO1* lines were used for measuring Ca^2+^ responses as described previously(Ueno et al., 2013). *UAS-synapto-pHluorin* (*UAS-spH*)(Ng et al., 2002) and *LexAop-synapto-pHluorin* (*LexAop-spH*) lines were used for measuring vesicle release(Ueno et al., 2013). *MB-LexA::GAD* (Ueno et al., 2013) was used for the *LexA* MB driver, *c747* (Aso et al., 2009) was used as the *GAL4* MB driver, and (Friggi-Grelin et al., 2003) and *TH-LexAp65* (Ueno et al., 2013) were used for *TH-DA* drivers. *UAS-shi^ts^* (Kitamoto, 2001) and *pJFRC104-13XLexAop2-IVS-Syn21-Shibire-ts1-p10* (*LexAop-shi^ts^*)(Pfeiffer et al., 2012) lines were used for inhibition of synaptic transmission. *MB247-Switch* (*MBsw*) was to express a *UAS* transgene in the MBs upon RU486 (RU+) feeding for 3-5 days(Mao et al., 2004). *UAS-dHO IR* was used to knockdown of *dHO* expression(Cui et al., 2008). *dHO*^Δ^ is a deficiency line *Df(3R)Exel7309* (BDSC 7960), lacking 65 Kbp including *dHO* (Flybase; http://flybase.org) in the third chromosome. *P{KK101716}VIE-260B* (VDRC ID 109631, Vienna Drosophila Resource Center, Vienna, Austria) (*UAS-RyR RNAi*) was used to knock down *RyR*. *Mi{Trojan-GAL4.0}RyR*[*MI08146-TG4.0*] (BDSC 67480) carries a *GAL4* sequence between 18 and 19 exon of *RyR* and was used to monitor *RyR* gene expression.

### Isolated whole brain preparation

Brains were prepared for imaging as previously described(Ueno et al., 2013). Briefly, brains were dissected in ice cold 0 mM Ca^2+^ HL3 medium (in mM, NaCl, 70; sucrose, 115; KCl, 5; MgCl_2_, 20; NaHCO_3_, 10; trehalose, 5; Hepes, 5; pH 7.3 and 359 mOsm)(Stewart et al., 1994), and placed in a recording chamber filled with normal, room temperature HL3 medium (the same recipe as above, containing 1.8 mM CaCl_2_). To deliver hemoCD through the blood brain barrier, brains were treated with papain (10 U/ml) for 15 min at room temperature, and washed several times with 0 mM Ca^2+^ HL3 medium prior to use(Gu and O’Dowd, 2007; Ueno et al., 2017).

### Imaging analysis

Imaging analysis was performed in HL3 solution as described previously (Ueno et al., 2013; Ueno et al., 2017). Briefly, fluorescent images were captured at 15 Hz using a confocal microscope system (A1R, Nikon Corp., Tokyo, Japan) with a 20x water-immersion lens (numerical aperture 0.5; Nikon Corp). We obtained *F_0_* by averaging the 5 sequential frames before stimulus onset and calculated Δ*F/F_0_*. To evaluate stimulation-induced fluorescent changes of spH, Δ*F/F_0_* calculated in the absence of stimulation or pharmacological agents was subtracted from stimulus or drug induced Δ*F/F_0._* To quantitatively evaluate the spH fluorescent changes, the average values of fluorescent changes at indicated time points during and after stimulation were statistically compared.

The AL was stimulated (30 pulses, 100 Hz, 1.0 ms pulse duration) using glass micro-electrodes. For NMDA stimulation, 200 μM NMDA, diluted in HL3 containing 4 mM Mg^2+^ (Miyashita et al., 2012), was applied by micro pipette.

For application of CO-saturated HL3, CO or control N_2_ gas was dissolved in HL3 saline by bubbling. CO or N_2_ saturated solutions were immediately placed in glass pipettes and puffed onto the MB lobes for 1 min (pressure = 6 psi) using a Picospritzer III system (Parker Hannifin Corp., USA). While we first used thin tip micropipettes, approximately 5 micro m diameter (Fig. 2*A*), we also used larger tip micropipettes, approximately 15 micro m (Fig. 5*E*), to avoid clogging.

To measure SV exocytosis using FFN511, *MB-LexA:GAD, LexAop-R-GECO1* brains were incubated in 10 μM FFN511/HL3 for 30 min. To remove non-specific binding of the dye, FFN511 loaded brains were washed in 200 μM ADVASEP-7/HL3 for 15 min two times(Kay et al., 1999; Gubernator et al., 2009). To evaluate stimulation-induced release of FFN511, Δ*F/F_0_* in the absence of stimulation was subtracted from Δ*F/F_0_*

### Behaviors

Olfactory aversive memory: The procedure for measuring olfactory memory has been previously described(Tully and Quinn, 1985; Tamura et al., 2003). Briefly, two mildly aversive odors (3-octanol [OCT]) or 4-methylcyclohexanol [MCH]) were sequentially delivered to approximately 100 flies for 1 min with a 45 sec interval between each odor presentation. When flies were exposed to the first, CS^+^ odor (either OCT or MCH), they were also subjected to 1.5 sec pulses of 60 V DC electric shocks every five sec. To test olfactory memory, flies were placed at the choice point of a T-maze where they were allowed to choose either the CS+ or CS-odor for 1.5 min. Memory was calculated as a performance index (PI), such that a 50:50 distribution (no memory) yielded a performance index of zero and a 0:100 distribution away from the CS^+^ yielded a performance index of 100.

Odor and Shock avoidance. Peripheral control experiments including odor and shock reactivity assays were performed as previously described(Tully and Quinn, 1985) to measure sensitivity to odors and electrical shocks. Approximately 100 flies were placed at the choice point of a T maze where they had to choose between an odor (OCT or MCH) and mineral oil or between electrical shocks and non-shocked conditions. A preformance index was calculated as described above.

### Identification of dHO localization

To detect of dHO protein in fly brains, wild-type, w(CS)(Dura et al., 1993), and *dHO*^Δ^ flies were dissected and fixed in 4 % paraformaldehyde for 20 min at 4°C. Brains were incubated in PBS with 5 % FBS and 0.1 % Triton-X for 30 min at 4°C, and then in primary antibodies, 1:50 anti-HO (Cui et al., 2008) and 1:20 anti-Fas2 (1D4, Developmental Studies Hybridoma Bank, Iowa, USA) for 3 days at 4°C. After washing, brains were incubated with secondary antibodies, Alexa488-conjugated donkey anti-rat antibody (1:200) (Invitrogen, Carlsbad, USA) and Alexa555-conjugated donkey anti-mouse antibody (1:200) (Invitrogen) for 2 days at 4°C. Images were captured using an A1R confocal microscope (Nikon, Tokyo, Japan).

### Identification of RyR positive neurons (Trojan)

For *Mi{Trojan-GAL4.0}RyR*[*MI08146-TG4.0*]*/UAS-mCD8::GFP* imaging, heads were dissected in 4 % paraformaldehyde for 30 min at 4°C. Brains were incubated with primary antibodies, anti-GFP (1:400) (ab13970, Abcam, Cambridge, UK) and anti-tyrosine hydroxylase (#22941, Immunostar, Hudson, USA) in PBS with 10 % ImmunoBlock (DS Pharma Biomedical Co., Osaka, Japan) and 0.1 % Triton-X overnight at 4°C. After washing, brains were incubated with secondary antibodies, Alexa488-conjugated donkey anti-chick antibody (1:400) (Jackson ImmunoResearch, West Grove, USA) and Alexa555-conjugated donkey anti-mouse antibody (1:400) (Invitrogen) overnight at 4°C. Images were captured using an A1R confocal microscope (Nikon, Tokyo, Japan).

### Chemicals and treatments

RU486 (mifepristone), NMDA (N-methyl-D-aspartate), L-NAME (Nϖ-L-nitro arginine methyl ester), 1-octanol, 2-arachidonil glycerol, arachidonic acid and CrMP (chromium mesoporphyrin) and ADVASEP-7 were purchased from Sigma-Aldrich (Missouri, USA). Thapsigargin and tetrodotoxin (TTX) were purchased from Wako Pure Chemical Industries (Osaka, Japan). Dantrolen was purchased from Alomone labs (Jerusalem, Israel). BAPTA-AM and O,O’-Bis(2-aminoethyl)ethyleneglycol-N,N,N’,N’-tetraacetic acid (EGTA) were purchased from Dojindo lab (Kumamoto, Japan). FFN511 was purchased from Abcam (Cambridge, England). Papain was purchased from Worthington Biochemical Corporation (New Jersey, USA). RU486 was dissolved in ethanol, butaclamol was dissolved in DMSO, L-NAME and SCH23390 were dissolved in water, CrMP was dissolved in 0.5% 2-aminoethanol and 2 mM HCl. Oxy-hemoCD was reduced in sodium dithionite and purified using a HiTrap Desalting column (GE Healthcare Japan, Tokyo, Japan) and eluted in PBS. The concentration of purified oxy-hemoCD is estimated by absorbance at 422 nm(Kitagishi et al., 2010). COP-1 and CORM-3 were prepared according to previous publications(Clark et al., 2003; Michel et al., 2012). Both reagents were stored at −20 °C and dissolved in DMSO before use. For RU486 treatment, dissolved RU486 was mixed in fly food. Flies were fed RU486 for 5 days prior to brain preparation. Other chemicals were treated as described in the main text and figure legends.

### Statistics

Statistical analyses were performed using Prism software (GraphPad Software, Inc., La Jolla, CA, USA). All data in bar and line graphs are expressed as means ± SEMs. Student’s t-test and Mann Whitney test was used to evaluate the statistical significance between two data sets. For multiple comparisons, one-way or two-way ANOVA followed by Bonferroni post hoc analyses were employed. Statistical significances are shown as \**P* < 0.05, \*\**P* < 0.01. *P* values greater than 0.05 were considered not statistically significant, NS > 0.05.

## RESULTS

### CO synthesis in the MB neurons is required for DA release upon coincident stimulation

Previously, we used an *ex vivo* dissected brain system to examine SV exocytosis from DA terminals projecting onto the α3/α’3 compartments of the vertical MB lobes. We measured SV exocytosis from DA terminals using a vesicular exocytosis sensor, synapto-pHluorin (spH)(Miesenbock et al., 1998), expressed in dopaminergic neurons using a tyrosine hydroxylase (TH) driver, and found that release occurred only upon coincident activation of post-synaptic MB neurons by cholinergic inputs from the ALs and glutamatergic inputs from the AFV(Ueno et al., 2017).

If postsynaptic MB activity evokes SV exocytosis from presynaptic DA terminals, vesicular output from MB neurons may be needed to activate DA neurons that loop back to the MBs, as has been previously suggested(Ichinose et al., 2015; Cervantes-Sandoval et al., 2017; Takemura et al., 2017; Horiuchi, 2019). To test this possibility, we inhibited synaptic transmission from MB neurons by expressing temperature-sensitive *shi^ts^* from a pan-MB driver, *MB-LexA*. We confirmed that MB synaptic output is blocked at restrictive temperature in *MB-LexA>LexAop-shi^ts^* flies by demonstrating that memory recall, which requires MB output(Dubnau et al., 2001; McGuire et al., 2001), is defective in these flies (Suppelemental Fig. S1*A*). Interestingly, SV exocytosis from TH-DA terminals occurs normally at restrictive temperature in these flies upon coincident AL + NMDA stimulation (Fig. 1*A*), suggesting while looping activity may be necessary for memory, it is not required for DA release. SV exocytosis from TH-DA terminals also occurred normally when *shi^ts^* was expressed using a different MB driver (*c747-GAL4>UAS-shi^ts^*) (Suppelemental Fig. S1*B*), even though memory recall was also disrupted at restrictive temperature in this line(Dubnau et al., 2001). 1 mM 1-octanol, a blocker of gap junctional communication(Rorig et al., 1996; Goncharenko et al., 2014), also did not inhibit SV exocytosis in TH-DA terminals (Fig. 1*B*). These results suggest that output from chemical and electrical synapses is not required for post-synaptic MB neurons to induce pre-synaptic DA-release from DA neurons.

**Fig. 1.**
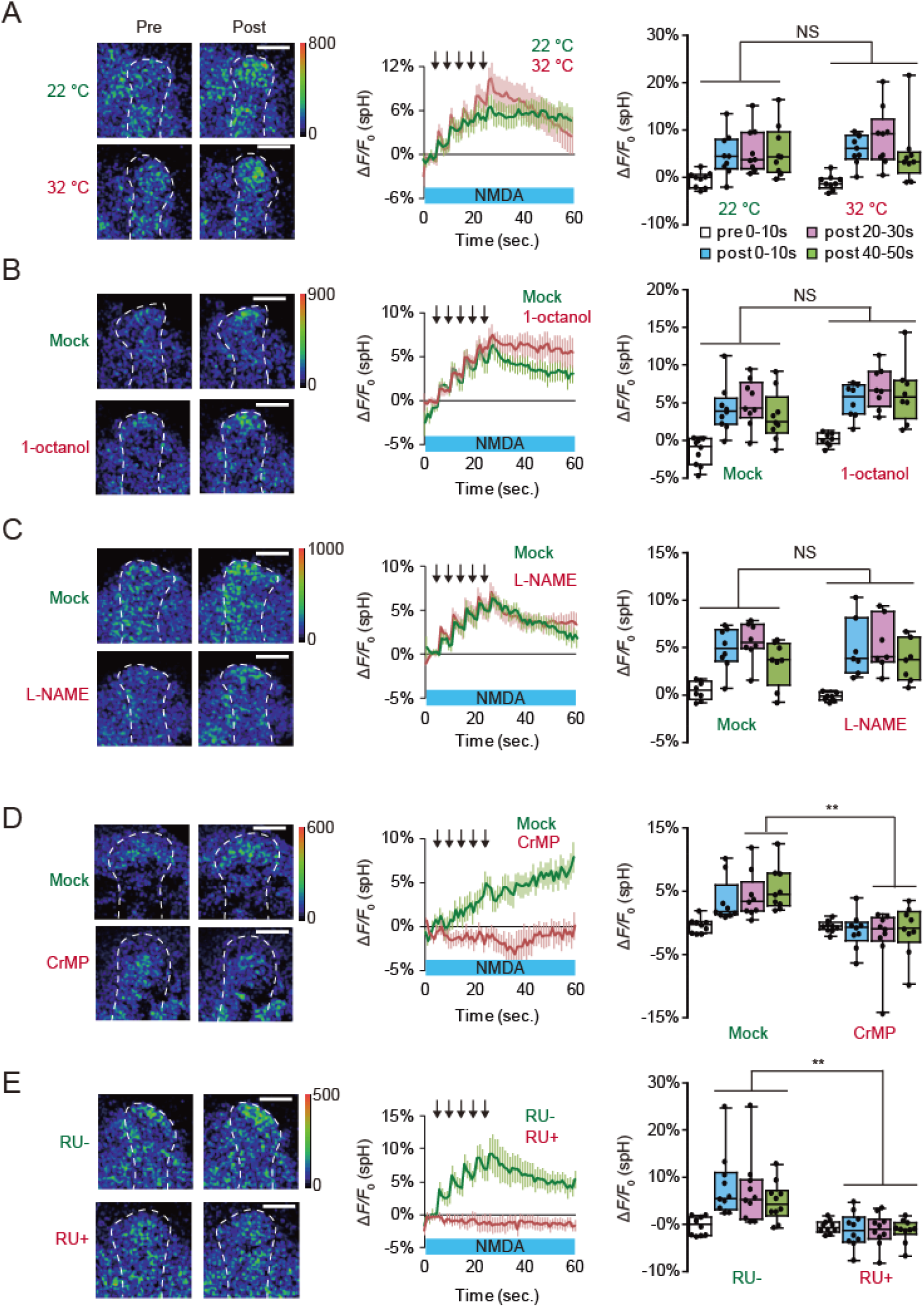
Inhibiting heme oxygenase (HO) activity in the MBs blocks SV exocytosis from pre-synaptic DA terminals. **A**, Blocking SV exocytosis from the MBs does not affect DA release. spH fluorescence was measured at TH-DA terminals innervating the α3/α’3 compartments of the MB vertical lobes in *MB-LexA>LexAop-shi^ts^*brains. (Left) typical pseudocolor images 5 s before and 30 s after coincident AL + NMDA activation. (Middle) time course of fluorescent changes, and (right) summary graphs. Two-way ANOVA indicates no significant differences in spH fluorescence between restrictive (32 °C) and permissive (22 °C) temperatures. N = 9 for all data. Scale bars = 20 μm. **B**, The gap junction inhibitor, 1-octanol, does not affect DA release. Two-way ANOVA indicates no significant differences in spH fluorescence between mock and 1-octanol conditions. N = 8 for all data. **C,** The NOS inhibitor, L-NAME, does not affect SV exocytosis from TH-DA terminals. Two-way ANOVA indicates no significant differences in spH fluorescence due to drug treatment. N=8 for all data. **D,** The HO inhibitor, CrMP, prevents DA release. Two-way ANOVA indicates significant differences in spH fluorescence due to drug treatment (*F_3, 64_* = 3.268, *P* = 0.0268, N = 9 for mock and N = 10 for 10 μM CrMP). * *P* < 0.05 and ** *P <* 0.01 by Bonferroni *post hoc* tests. **E**, *dHO* knockdown in the MBs prevents DA release. Two-way ANOVA indicates significant differences in spH fluorescence due to RU treatment (*F_3, 72_* = 4.265, *P* = 0.0079, N = 10 for all data). ** *P <* 0.01 by Bonferroni *post hoc* tests.

We next examined whether a retrograde messenger, such as nitric oxide (NO), may be released from MB neurons to regulate pre-synaptic DA release. However, 100 μM L-NAME (Nϖ-L-nitro arginine methyl ester), a NO synthetase blocker(Boultadakis and Pitsikas, 2010) had no effect on AL + NMDA stimulation-induced SV exocytosis from TH-DA terminals (Fig. 1*C*).

Olfactory memory is disrupted by mutations in *nemy*, a gene that encodes a *Drosophila* homolog of cytochrome B561 (CytB561)(Iliadi et al., 2008), which is involved in metabolism of carbon monoxide (CO)(Sugimura et al., 1980; Cypionka and Meyer, 1983; Jacobitz and Meyer, 1989), a diffusible gas similar to NO, that also been proposed to act as a retrograde messenger during synaptic plasticity(Alkadhi et al., 2001; Shibuki et al., 2001). Thus, we next examined whether CO may be required for DA release. CO is synthesized by heme oxygenase (HO), and we found that exocytosis from TH-DA terminals upon coincident activation of MB neurons is abolished upon application of chromium mesoporphyrin (CrMP), a HO blocker(Vreman et al., 1993) (Fig. 1*D*). LTE was also significantly inhibited by CrMP (Suppelemental Fig. S2*A*), demonstrating the importance of DA release in plasticity. To verify that the *Drosophila* homologue of HO (dHO)(Cui et al., 2008) is present in the MBs, we used anti-dHO antibodies and found strong expression in the MBs and in insulin producing cells (Suppelemental Fig. S2*B*). We next inhibited *dHO* expression in the MBs using *MBsw>UAS-dHO-IR* flies, which express a *dHO-RNAi* construct from an RU486-inducible *MB247-switch* (*MBsw*) driver(Mao et al., 2004). We found that both SV exocytosis from TH-DA terminals (Fig. 1*E* and Suppelemental Movie S1*A, B*) as well as LTE (Suppelemental Fig. S2*C*) were impaired when these flies were fed RU486. Furthermore, acute knock down of dHO in the MBs in *MBsw>UAS-dHO-IR* flies disrupted olfactory memory (Suppelemental Fig. S2*D*) without affecting task-related responses (Suppelemental Fig. S2*E*). Altogether, these results indicate that dHO in the MBs is required for olfactory memory, MB plasticity, and DA release onto MBs.

### CO generated from coincidentally activated MB neurons evokes DA release

If CO functions as a retrograde messenger inducing DA release, direct application of CO to DA terminals should induce release. Thus we next applied CO-saturated saline from micropipettes to the vertical lobes of the MBs, and observed robust SV exocytosis from TH-DA terminals (Fig. 2*B* and Suppelemental Movie S2). Further, we found that application of a CO-releasing molecule-3 (CORM-3), a water-soluble CO-donor(Tinajero-Trejo et al., 2014; Aki et al., 2018), also evokes SV exocytosis from TH-DA terminals (Fig. 2*B*). In contrast, application of other retrograde messengers, including 200 μM arachidonic acid and 200 μM 2-arachidonylglycerol, an endocanabinoid receptor agonist, had no effects on release (Fig. 2*C* and 2*D*). To further examine whether endogenously generated CO is required for DA release, we used a CO selective scavenger, hemoCD(Kitagishi et al., 2010) and found that hemoCD significantly inhibited vesicular exocytosis from TH-DA terminals upon AL + NMDA stimulation (Fig. 2*E*).

**Fig. 2.**
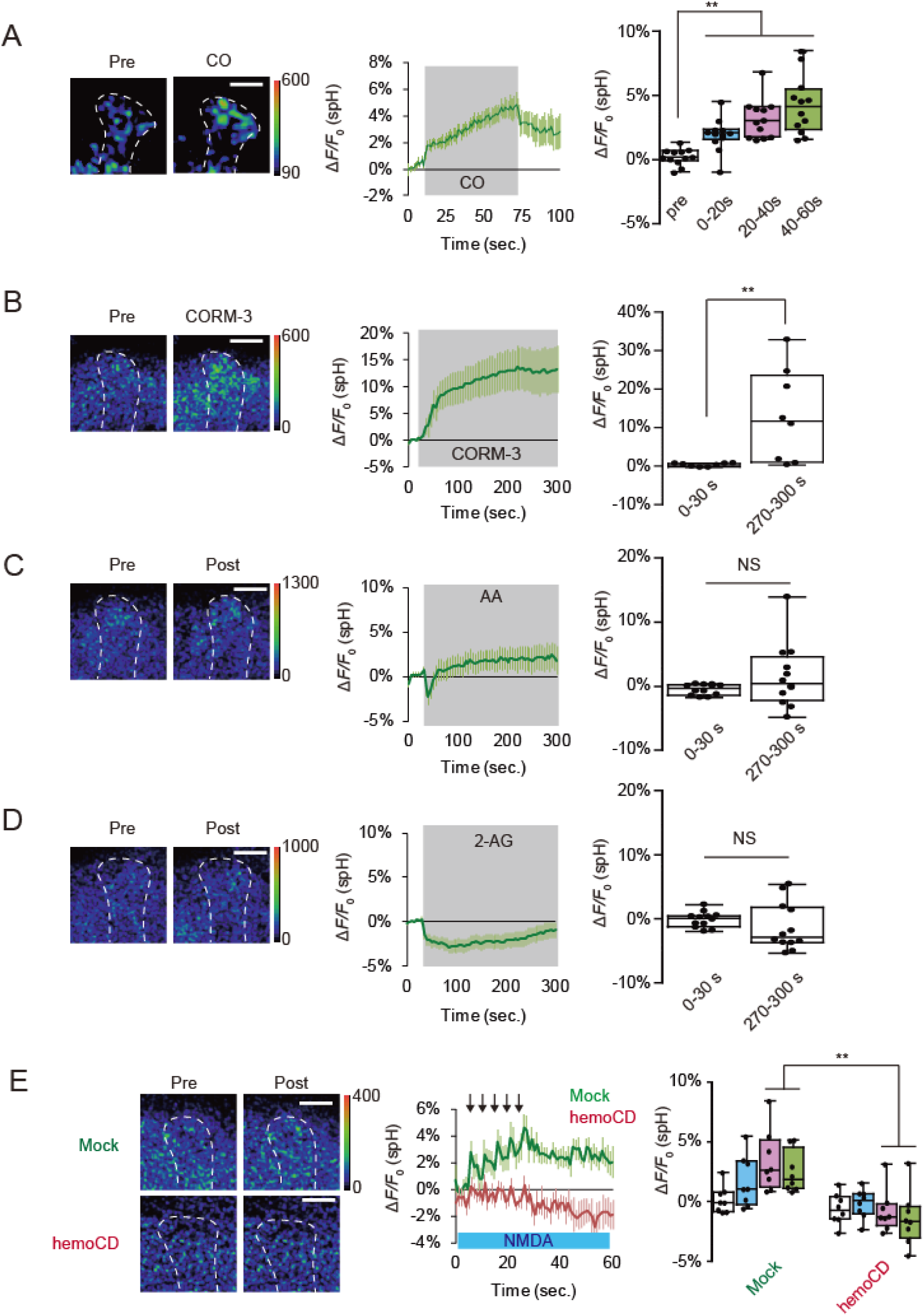
CO evokes SV exocytosis from DA terminals. **A**, CO induces DA release. CO-saturated saline was puffed onto the vertical lobes of the MB during the indicated time period (gray-shaded area in the middle panel). CO-induced spH responses were obtained by subtracting spH fluorescent changes induced by control N_2_-satulated saline from fluorescence changes induced by CO-saturated saline. One-way ANOVA and post hoc tests indicate significant changes in spH fluorescence upon CO application (*F_3,44_*= 14.994, *P* < 0.001). N = 12 for all data. **B**, CORM-3 induces DA release. 20 μM CORM-3 (dissolved in 0.2% DMSO) was bath applied and spH fluorescence was measured at DA terminals at the tip of the MB vertical lobes. CORM-3-induced spH responses were obtained by subtracting fluorescent changes induced by control 0.2% DMSO saline from fluorescence changes induced by CORM-3. ** *P* < 0.01 determined by Student’s t-test, comparing spH responses before (0-30s) and after application of CO (270-300s). N = 8 for all data. **C**, Arachidonic acid (AA) does not induce DA release. 200 μM AA (dissolved in 0.4 % ethanol) was bath applied. AA-induced spH responses were obtained by subtracting fluorescence changes induced by 0.4% ethanol saline from fluorescence changes induced by AA. NS *P* > 0.05 by Mann Whitney test and N = 16 for all data. **D,** Effects of 2-arachidonylglycerol (2-AG) on DA release. Bath application of 200 μM AG (dissolved in 0.2 % DMSO) did not induce SV exocytosis in TH-DA terminals. AG-induced spH responses were obtained by subtracting fluorescence changes induced by 0.2% DMSO saline from fluorescence changes induced by AG. NS *P* > 0.05 determined by Student’s t-test. N = 12 for all data. **E,** The CO scavenger, hemoCD, prevents SV exocytosis from DA terminals. AL + NMDA-dependent changes in spH fluorescence were measured in the presence and absence of 50 μM hemoCD. Two-way ANOVA indicates significant decreases in spH fluorescence due to hemoCD treatment (*F_3, 56_*= 2.845, *P* = 0.0458, N = 8 for all data). ** *P <* 0.01 by Bonferroni *post hoc* tests.

We next visualized the generation and release of CO from MB neurons using a CO selective fluorescent probe, CO Probe 1 (COP-1)(Michel et al., 2012). While COP-1 fluorescence increased immediately after coincident AL + NMDA stimulation, fluorescence remained unchanged after AL stimulation or NMDA application alone (Fig. 3*A*). Thus, changes in COP-1 fluorescence parallel changes in DA release. Furthermore, the fluorescence increase in COP-1 occurred on the lobes of the MBs ipsilateral, but not contralateral, to the stimulated AL (Fig. 3*B* and Suppelemental Movie S3). Since each AL innervates its ipsilateral, but not contralateral MB, this suggests that CO production occurs in areas of coincident AL and NMDA activation. Again this result parallels that of DA release(Ueno et al., 2017). Significantly, increased COP-1 fluorescence was attenuated by knocking down dHO in the MBs (Fig. 3*C*), indicating that COP-1 flurorescence detects dHO-dependent CO production. Collectively, these results suggest that coincident stimulation of MB neurons induces dHO to generate the retrograde messenger, CO, which then evokes SV exocytosis from presynaptic DA terminals.

**Fig. 3.**
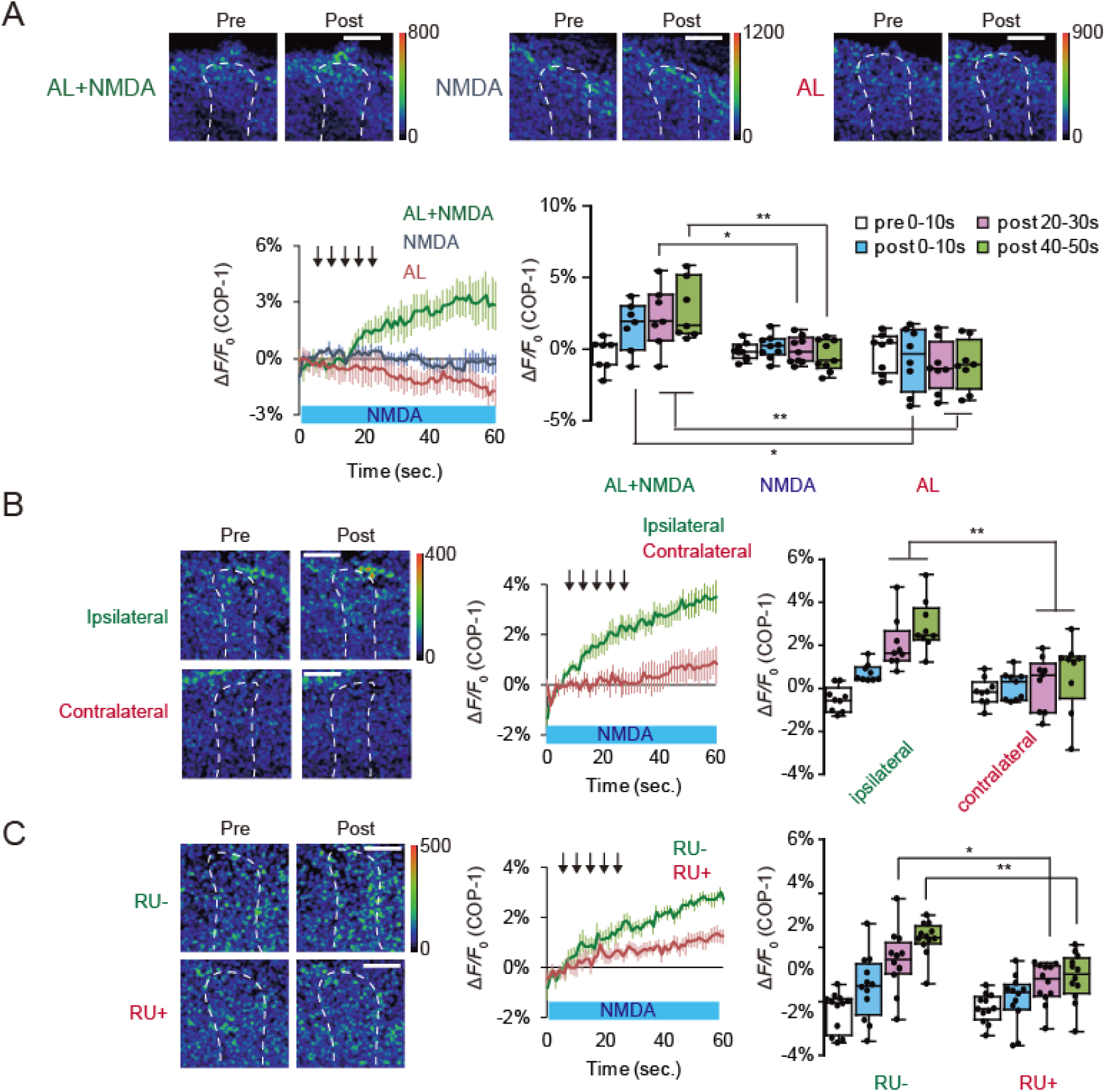
Endogenous CO released from MBs induces DA release. **A**, Top panels, typical images of COP-1 fluorescence observed at the α3/α’3 compartments of the MB vertical lobes. Pre refers to images taken prior to stimulation, while Post refers to images taken 30 secs after onset of indicated stimuli. Lower left, time course of COP-1 fluorescence upon indicated stimulation protocols, and lower right, summary of COP-1 fluorescence at different time intervals upon indicated stimulation. Isolated brains were incubated in 4 μM COP-1, and COP-1 responses were calculated by subtracting non-stimulated fluorescence changes from stimulated fluorescence changes. Two-way ANOVA indicates significant differences in COP-1 fluorescence due to treatment, time, and interaction between treatment and time (*F_6, 84_ =* 3.094, *P =* 0.0087, N = 8 for AL or NMDA stimulation alone and N = 7 for AL + NMDA stimulation). * *P* < 0.05 and ** *P <* 0.01 by Bonferroni *post hoc* tests. Scale bars = 20 μm. **B,** CO is released from the MB lobe ipsilateral to AL stimulation. Two-way ANOVA indicates significant differences in fluorescence between MB lobes (*F_3, 64_ =* 5.491, *P =* 0.020, N = 9 for all data). ** *P <* 0.01 by Bonferroni *post hoc* tests. **C,** Knocking down *dHO* expression in the MBs impairs CO production. Two-way ANOVA indicates significant differences in COP-1 fluorescence due to RU treatment (*F_3, 88_* = 6.25, *P =* 0.0086, N = 12 for all data). ** *P <* 0.01 by Bonferroni *post hoc* tests.

### CO evokes non-canonical SV exocytosis

SV exocytosis requires an increase in Ca^2+^ concentration in presynaptic terminals(Katz and Miledi, 1967; Augustine et al., 1985; Sabatini and Regehr, 1996; Meinrenken et al., 2003). Consistent with this, we observed a robust Ca^2+^ increase in TH-DA terminals that project onto the MB lobes receiving coincident AL + NMDA stimulation (Fig. 4*A* and Suppelemental Movie S4), but not in terminals that project to the contralateral side (Fig. 4*B*). Ca^2+^ increases in DA terminals were also observed upon application of CORM-3 (Fig. 4*C*), and this increase was abolished by addition of the membrane-permeable Ca^2+^ chelator BAPTA-AM (Fig. 4*D*). In canonical SV exocytosis, neuronal activity opens voltage-gated calcium channels, allowing influx of extracellular Ca^2+^(Katz and Miledi, 1967; Augustine et al., 1985). However, we found that CORM-3 is able to induce SV exocytosis from TH-DA terminals even in Ca^2+^ free saline in the presence of the Ca^2+^ chelator, EGTA, and the sodium channel blocker, TTX (Fig. 4*E*). This suggests that CO-evoked DA release is action potential independent and does not require Ca^2+^ influx from extracellular sources.

**Fig. 4.**
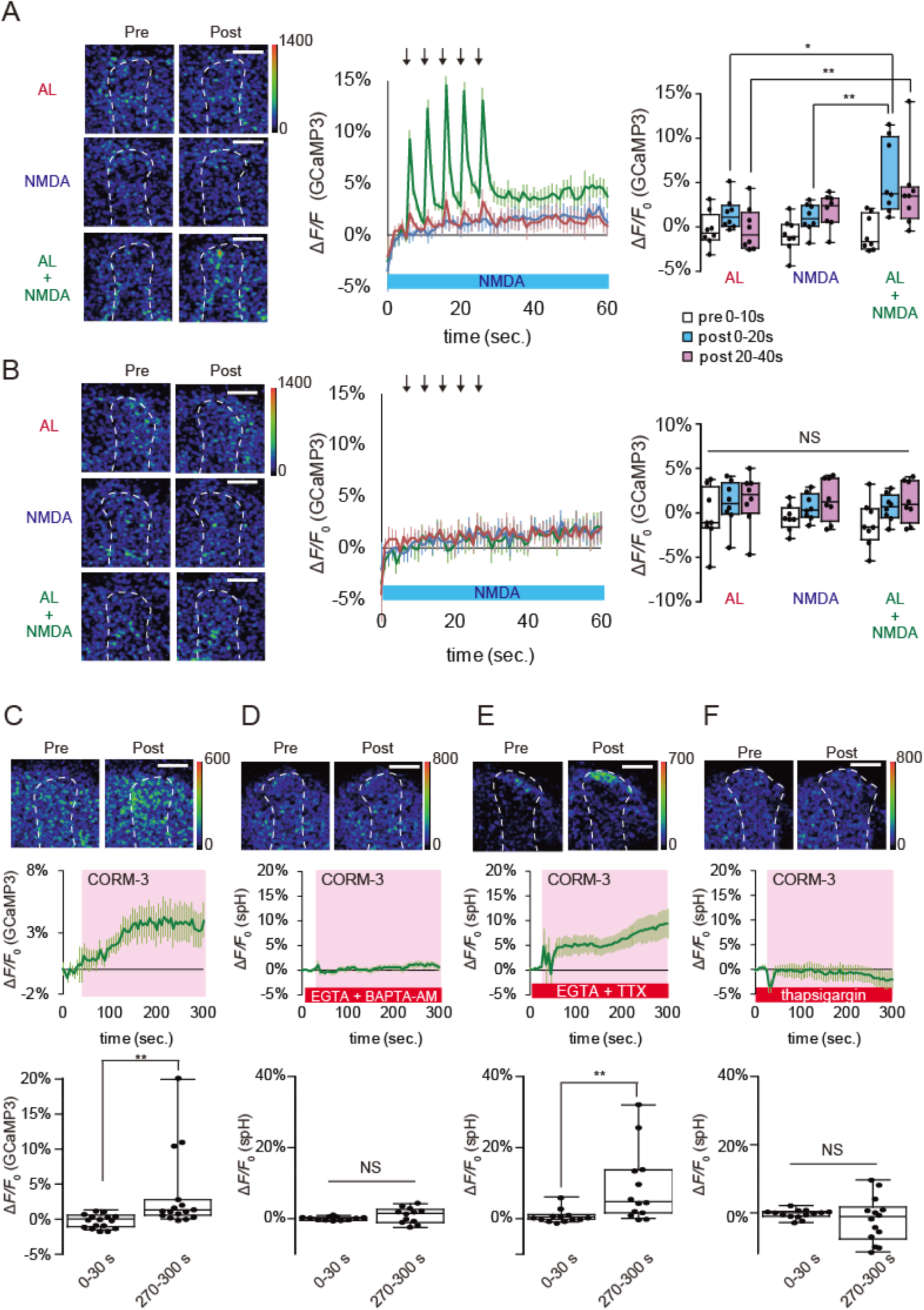
CO induced DA release requires Ca^2+^ efflux from internal Ca^2+^ stores. **A**, Left panel, typical pseudo color images of G-CaMP3 fluorescence in TH-DA terminals innervating the α3/α’3 compartments of the vertical MB lobes ipsilateral to AL stimulation. The type of stimulation applied is indicated on the left, and responses before (pre) and after (post) stimulation are shown. Middle panel, time course of Ca^2+^ responses in TH-DA terminals upon indicated stimulation. Right panel, summary of responses. Two-way ANOVA indicates significant differences in G-CaMP3 fluorescence due to stimulation type, time, and interaction between stimulation type and time (*F_4, 63_* = 2.610, *P =* 0.0437, N = 8 for all data). * *P* < 0.05 and ** *P <* 0.01 by Bonferroni *post hoc* tests. **B**, G-CaMP3 fluorescence changes in TH-DA terminals innervating the MB contralateral to AL stimulation. Panels are similar to those shown in (A). Two-way ANOVA indicates no significant differences in G-CaMP3 fluorescence due to time or stimulation type. N = 9 for all data. **C**, CORM-3 induces Ca^2+^ increases in TH-DA terminals. CORM-3 induced Ca^2+^ changes were calculated by subtracting fluorescence changes under mock conditions from changes induced by CORM-3. 20 μM CORM-3 (dissolved in 0.2% DMSO saline) was puffed onto the α3/α’3 compartments of the vertical MB lobes during the indicated time period (shown in pink in the middle panel). \*\**P* < 0.01 by Mann Whitney test. N = 16 for all data. **D**, CORM-3-induced DA release depends on increases in intracellular Ca^2+^. Brains were incubated in Ca^2+^-free external solution containing 2 mM EGTA and 10 μM BAPTA-AM for 10 min prior to CORM-3 application. *P* = 0.166 by Student’s t-test. N = 12. **E**, CORM-3-induced DA release does not require influx of external Ca^2+^ or activity of voltage-gated Na^+^ channels. Brains were placed in Ca^2+^-free external solution containing 5 mM EGTA and 10 μM TTX 10 min prior to CORM-3 application. \*\**P* < 0.01 by Mann Whitney test. N = 13. **F**, CORM-3-induced DA release requires efflux from internal Ca^2+^ stores. Brains were incubated in Ca^2+^-free external solution containing 10 μM thapsigargin for 10 min prior to CORM-3 application. *P* = 0.364 by Student’s t-test. N = 14.

Since extracellular Ca^2+^ is not responsible for CO-dependent DA release, we next examined whether Ca^2+^ efflux from internal stores may be required. Significantly, CORM-3 failed to increase Ca^2+^ in TH-DA terminals in the presence of EGTA and thapsigargin, an inhibitor of the sarcoplasmic/endoplasmic reticulum Ca^2+^ ATPase (SERCA), which depletes internal Ca^2+^ stores(Kijima et al., 1991; Sagara and Inesi, 1991) (Fig. 4*F*). Thus, CO-evoked DA release occurs through a non-canonical mechanism that depends on Ca^2+^ efflux from internal stores rather than from extracellular sources.

### Ryanodine receptors mediate Ca^2+^ efflux for CO-evoked DA release

What mediates Ca^2+^ efflux from internal stores in DA terminals? Inositol 1,4,5-trisphosphate receptors (IP_3_Rs) and ryanodine receptors (RyRs) are the major channels mediating Ca^2+^ release from internal stores(Bardo et al., 2006). While SV exocytosis evoked by coincident MB stimulation was not suppressed by 2-Aminoethoxydiphenyl borate (2-APB), an IP_3_R antagonist(Maruyama et al., 1997) (Fig. 5*A*), exocytosis was significantly inhibited by dantrolene, a RyR antagonist(Zhao et al., 2001) (Fig. 5*B*). Conversely, application of a RyR agonist, 4-chloro-3-methylphenol (4C3MP)(Zorzato et al., 1993) was sufficient to evoke exocytosis (Fig. 5*C*). These data suggest that RyRs in DA neurons are required for SV exocytosis upon coincident activation of MB neurons.

**Fig. 5.**
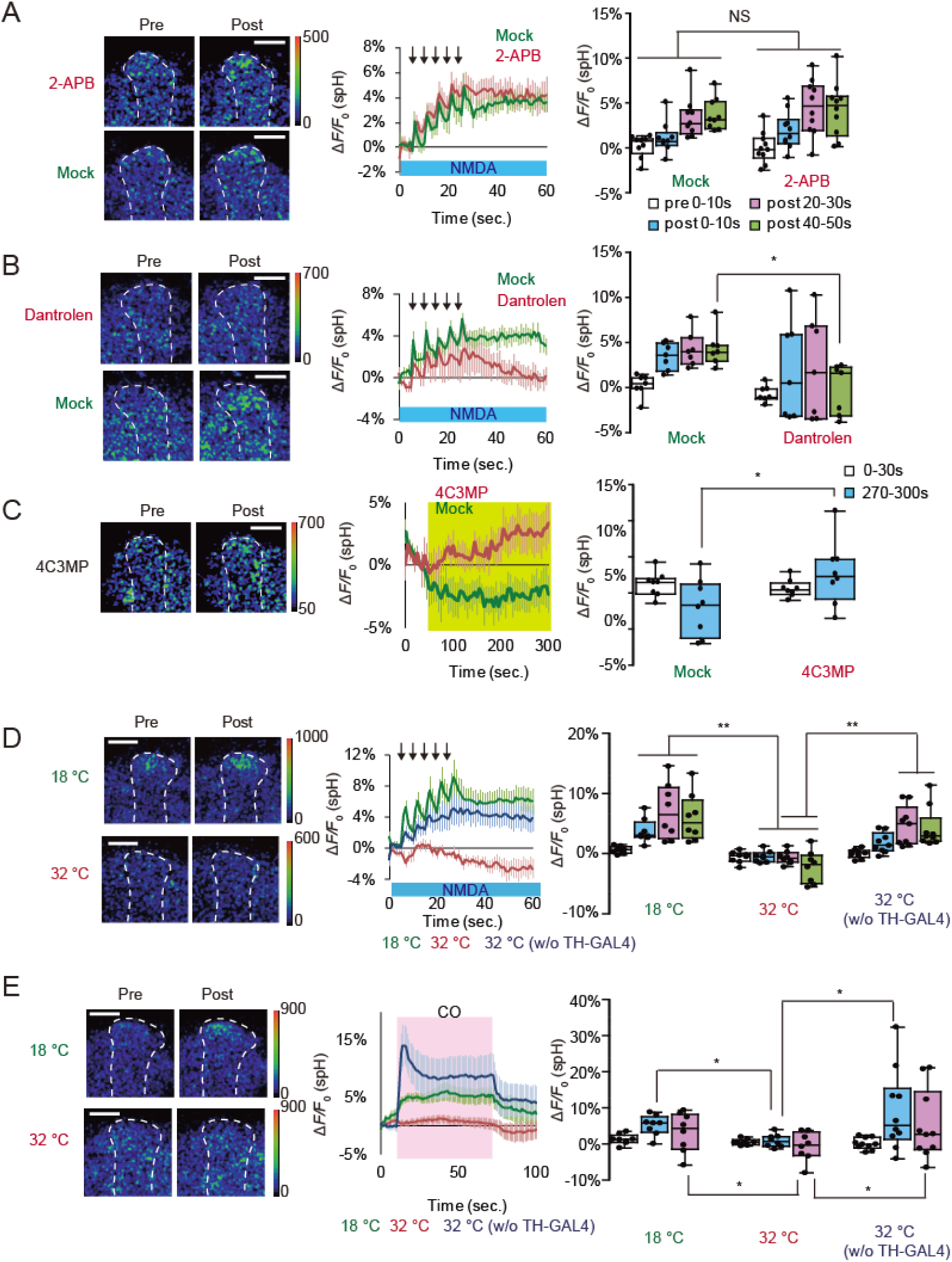
CO induced DA release is mediated by ryanodine receptors. **A**, An IP_3_R inhibitor does not affect DA release. 100 μM 2-aminoethoxydiphenylborane (2-APB) was dissolved in 0.1 % DMSO saline. Two-way ANOVA indicates no significant differences in spH fluorescence due to 2-APB treatment. N = 10 for 2-APB treatment and N = 9 for mock control. **B**, The RyR inhibitor, dantrolen, inhibits DA release. 10 μM dantrolen was dissolved in 0.1% DMSO saline. Two-way ANOVA indicates significant effects of drug treatment on spH fluorescence (*F_1, 48_* = 6.781, *P* = 0.0122, N = 8 for all data). \**P* < 0.05 compared with mock treated samples assayed by Bonferroni *post hoc* tests. **C**, Application of a RyR agonist induces DA release. 1 mM 4-chloro-3-methylphenol (4C3MP) was dissolved in 0.2% ethanol saline containing 1 μM TTX and applied for the indicated period of time (shown in yellow). ** *P* < 0.001 as assayed by Student’s t-test comparing before (0-30 s) and after (280-300 s) 4C3MP treatment. N = 8 for all data. **D**, Temporal *RyR* knockdown at 30 °C in TH-DA neurons prevents DA release evoked by AL + NMDA stimulation. Two-way ANOVA indicates signifincant differences in spH fluorescence between restrictive and permissive temperatures (*F_3,56_*= 6.625, *P* = 0.0007, N = 8 for all data). ** *P <* 0.01 by Bonferroni *post hoc* tests. **E**, Temporal knock down of *RyR* in TH-DA neurons prevents DA release evoked by CO application. CO saturated saline was applied during the indicated time period (shown in pink) from a micropipet. Two-way ANOVA indicates significant interaction differences in spH fluorescence due to time, temperature, and interaction between time and temperature (*F_2,42_* = 12.6, *P* < 0.0001, N = 8 for all data). ** *P <* 0.01 by Bonferroni *post hoc* tests.

To address whether RyRs are expressed in DA terminals, we examined expression of mCD8::GFP in *Mi{Trojan-GAL4.0}RyR^MI08146-TG4.0^; P{UAS-mCD8::GFP}* flies. In *Mi{Trojan-GAL4.0}RyR^MI08146-TG4.0^*, a Trojan GAL4 exon is inserted between exons 18 and 19 in the same orientation as the RyR gene(Diao et al., 2015). Thus GAL4, and mCD8::GFP, expression should reflect RyR expression. In these flies, mCD8::GFP signals overlapped with anti-TH antibody signals, indicating that RyRs are expressed in DA neurons (Suppelemental Fig. 3*A*). To determine whether RyRs in the DA terminals are required for DA release, we used the TARGET system(McGuire et al., 2003) to acutely knock down RyRs in adult TH-DA neurons, and found that this significantly suppressed SV exocytosis from DA terminals upon AL + NMDA stimulation (Fig. 5*D*). Furthermore, acute knock down RyRs also suppressed SV exocytosis induced by direct CO application to TH-DA terminals (Fig. 5*E*), and also suppressed LTE upon coincident AL + NMDA stimulation (Suppelemental Fig. S3*B*). Thus, pre-synaptic RyRs are required for both activation-dependent and CO-dependent DA release, MB plasticity, and olfactory memory.

## DISCUSSION

### CO functions as a retrograde on-demand messenger for SV exocytosis in presynaptic DA terminals

A central tenet of neurobiology is that action potentials, propagating from the cell bodies, induce Ca^2+^ influx in presynaptic terminals to evoke SV exocytosis. However, recent mammalian studies have shown that only a certain fraction of a large number of pre-synaptic release sites is involved in canonical SV exocytosis(Pereira et al., 2016; Liu et al., 2018). In this study we identify a novel mechanism of SV exocytosis in which activity in post-synaptic neurons evokes pre-synaptic release to induce plastic changes. This mechanism allows the timing and location of DA release to be strictly defined by activity of postsynaptic neurons.

On-demand SV exocytosis utilizes CO as a retrograde signal from postsynaptic MB neurons to presynaptic DA terminals. We demonstrate that CO fulfills the criteria that have been proposed for a retrograde messenger(Regehr et al., 2009). First, we demonstrate that HO, which catalyzes CO production, is highly expressed in postsynaptic MB neurons, indicating that MB neurons have the capacity to synthesize the messenger. Second, we show that pharmacological and genetic suppression of HO activity in the MBs inhibits CO production, pre-synaptic DA release, and LTE. Third, using a CO fluorescent probe, COP-1(Michel et al., 2012), we demonstrate that CO is generated in the MBs following coincident stimulation of the MBs, and CO generation is restricted to lobes of MB neurons that receive coincident stimulation. We further show that direct application of CO, or a CO donor, induces DA release from presynaptic terminals, while addition of a CO scavenger, HemoCD, suppresses release. Fourth, we demonstrate that CO activates RyRs in presynaptic terminals to induce SV exocytosis. Strikingly, CO-dependent SV exocytosis does not depend on influx of extracellular Ca^2+^, but instead requires efflux of Ca^2+^ from internal stores via RyRs. Finally, we show that pharmacological inhibition and genetic suppression of RyRs in DA neurons impairs DA release after coincident stimulation and CO application.

Other retrograde signals, such as NO and endo cannabinoids enhance or suppress canonical SV exocytosis, we find that CO-dependent DA release occurs even in conditions which block neuronal activity and Ca^2+^ influx in presynaptic DA terminals. This suggests that CO does not function to modulate canonical SV exocytosis, but may instead evoke exocytosis through a novel mechanism. Several previous studies have indicated that CO and RyR-dependent DA release also occurs in mammals. A microdialysis study has shown that CO increases the extracellular DA concentration in the rat striatum and hippocampus(Hiramatsu et al., 1994), either through increased DA release, or inhibition of DA reuptake(Taskiran et al., 2003). Also, pharmacological stimulation of RyRs has been reported to induce DA release in the mice striatum(Oyamada et al., 1998; Wan et al., 1999). This release is attenuated in RyR3 deficient mice, while KCl-induced DA release, which requires influx of extracellular Ca^2+^, is unaffected, suggesting that RyR-dependent release is distinct from canonical DA release. However, it has been unknown whether and how CO is generated endogenously. Also physiological conditions that activate RyRs to evoke DA release have also been unclear.

While our results demonstrate that CO signaling is necessary and sufficient for DA release, we note that our studies use fluorescent reporters which are not optimal for detailed kinetic analysis of release and reuptake. For example, increases in spH fluorescence can be used to determine vesicular release from DA neurons, but the decrease in spH fluorescence after release does not reflect the kinetics of clearance of DA from synaptic sites. Similarly, increases in COP-1 fluorescence reflect increases in CO production and release, but the kinetics of this increase depends on CO binding affinities and limit of detection (Suppelemental Fig. S*4*) as well as CO production, and the gradual increase in COP-1 fluorescence following coincident activation does not indicate that CO production is similarly gradual. In addition, since COP-1 binding to CO is irreversible, we do not see a decrease in COP-1 fluorescence after the end of stimulation. Thus, although our functional imaging studies reliably measure significant changes in synaptic release, calcium signaling, and CO production, they are not precise enough to accurately measure the fast dynamics of these changes.

### Signaling pathway for CO-dependent on-demand release of DA

While most neurotransmitters are stored in synaptic vesicles and released upon neuronal depolarization, the release of gaseous retrograde messengers such as NO and CO is likely coupled to activation of their biosynthetic enzymes, NOS and HO. Previously, we demonstrated that activity-dependent DA release onto the MBs requires *rutabaga* adenylyl cyclase (*rut*-AC) in the MBs(Ueno et al., 2017). *rut*-AC is proposed to function as a neuronal coincidence detector that senses coincident sensory inputs and activates the PKA pathway by increasing production of cAMP. In mammals, activation of the cAMP/PKA pathway increases expression of an HO isoform, HO-1, through transcriptional activation of the transcription factor CREB(Durante et al., 1997; Park et al., 2013; Astort et al., 2016). However, gene expression changes are not fast enough to explain CO-dependent DA release. A second mammalian HO isoform, HO-2 is selectively enriched in neurons, and HO-2-derived CO is reported to function in plasticity. HO-2 is activated by Ca^2+^/calmodulin (CaM) binding(Boehning et al., 2004), and by casein kinase II (CKII) phosphorylation(Boehning et al., 2003), two mechanisms that can generate CO with sufficient speed to account for LTE. Currently, it is unclear whether *rut*-AC and the cAMP/PKA pathway functions in parallel with Ca^2+^/CaM and CKII, or in concert with these pathways (ie functions as a priming kinase for CKII(Huang et al., 2007)) to activate HO. Supporting the concept that PKA regulates HO, in the golden hamster retina, PKA has been shown to increase HO activity without affecting HO gene expression(Sacca et al., 2003).

While *Drosophila* has a single isoform of RyR, mammals have three isoforms, RyR1 to RyR3. Skeletal muscle and cardiac muscle primarily express RyR1 and RyR2, and the brain, including the striatum, hippocampus and cortex, expresses all three isoforms(Giannini et al., 1995). RyRs are known to be activated by Ca^2+^ to mediate Ca^2+^ induced Ca^2+^ release (CICR)(Endo, 2009). However, CO-evoked DA release occurs even in the presence of Ca^2+^-free extracellular solutions containing TTX and EGTA, suggesting that CO activates RyRs through a different mechanism. Besides Ca^2+^, RyRs can be activated by calmodulin, ATP, PKA, PKG, cADP-ribose, and NO(Takasago et al., 1991; Xu et al., 1998; Verkhratsky, 2005; Zalk et al., 2007; Lanner et al., 2010; Kakizawa, 2013). NO can directly stimulate RyR1 non-enzymatically by S-nitrosylating a histidine residue to induce Ca^2+^ efflux(Xu et al., 1998; Kakizawa, 2013). Similarly, CO has been reported to activate Ca^2+^-activated potassium channels (K_Ca_) through a non-enzymatic reaction in rat artery smooth muscle(Wang and Wu, 2003). Alternatively, both NO and CO can bind to the heme moiety of soluble guanlylate cyclase (sGC) leading to its activation(Stone and Marletta, 1994). Activated sGC produces cGMP, and cGMP-dependent protein kinase (PKG) rapidly phosphorylates and activates RyRs(Takasago et al., 1991). Interestingly, NO increases DA in the mammalian striatum in a neural activity independent manner(Hanbauer et al., 1992; Zhu and Luo, 1992; Lonart et al., 1993). Since activation of RyRs also increases extracellular DA in the striatum, hippocampus and cortex(Oyamada et al., 1998; Wan et al., 1999), NO may play a pivotal role in RyR activation and DA release in mammals. However, NOS expression has not been detected in the MBs(Muller, 1994; Regulski and Tully, 1995), suggesting that in *Drosophila*, CO rather than NO may function in this process.

### Biological significance of on-demand DA release

DA plays a critical role in associative learning and synaptic plasticity(Huang and Kandel, 1995; Jay, 2003; Puig et al., 2014; Lee et al., 2016; Yamasaki and Takeuchi, 2017). In flies, neutral odors induce MB responses by activating sparse subsets of MB neurons. After being paired with electrical shocks during aversive olfactory conditioning, odors induce larger MB responses in certain areas of the MBs(Yu et al., 2006; Wang et al., 2008; Akalal et al., 2011; Davis, 2011). We modeled this plastic change in *ex vivo* brains as LTE, and showed that DA application alone is sufficient to induce this larger response(Ueno et al., 2017). However, in the *Drosophila* brain, only a small number of DA neurons (∼12 for aversive and ∼100 for appetitive) regulate plasticity in ∼2000 MB Kenyon cells(Mao and Davis, 2009). Thus to form odor-specific associations, there should be a mechanism regulating release at individual synapses. CO-dependent on-demand DA release provides this type of control. If on-demand release is involved in plasticity and associative learning, knockdown of genes associated with release should affect learning. Indeed, we show that knocking down either dHO in the MBs or RyRs in DA neurons impairs olfactory conditioning. While these knockdowns did not completely abolish olfactory conditioning, this may be due to inefficiency of our knockdown lines. Alternatively, on-demand release may not be the only mechanism responsible for memory formation, but may instead be required for a specific phase of olfactory memory.

In olfactory aversive conditioning, CS+ odor is paired with US electrical shock. Given that AFV conveys US information, it seems strange that DA is not released by AFV stimulation alone in our *ex vivo* imging, while previous *in vivo* imaging study demonstrated that DA is released by electrical shock presentation alone (Sun et al., 2018). Notably, projection of DA terminals are compartmentalized on the MB lobes and show distinct responses and DA release during sensory processing (Cohn et al., 2015; Sun et al., 2018). In our *ex vivo* imaging, we looked at DA release onto the α3/α’3 compartments of the MB vertical lobes, while the previous *in vivo* imaging study found DA release onto the γ2 and γ3 compartments of the MB horizontal lobes upon electrical shock (Sun et al., 2018). Therefore, AFV stimulation may also induce DA release onto these horizontal compartments although it does not induce DA release onto α3/α’3 compartments. However, due the location of the microelectrode for AL stimulation, we did not look at the horizontal lobes in this study. Another possibility is that arousal state of the flies in *in vivo imaging* may be essential for DA release upon shock presentation. While our *ex vivo* imaging study supposes that glutamatergic neurons transmit AFV information to the MBs (Ueno et al., 2017), aversive US information has been proposed to be transmitted by DA neurons (Claridge-Chang et al., 2009; Aso et al., 2010; Aso et al., 2012; Burke et al., 2012; Liu et al., 2012). Synaptic terminals immunoreactive for the vesicular glutamate transporter, VGLUT, have been identified at the α3 compartment of the MB vertical lobes (Daniels et al., 2008). However, recent connectome study demonstrates that they are presynaptic terminals of the MB output neurons (Takemura et al., 2017). Nevertheless, it is noteworthy that these immunohisitochemical and connectome data do not correlate with expression of NMDA receptors in the MBs (Miyashita et al., 2012). These mismatch observations implicate that AFV mediated shock information may be transmitted to the MBs by another type of glutamatergic neurons.

In mammals, the role of CO in synaptic plasticity is unclear. Application of CO paired with low frequency stimulation induces LTP, while inhibiting HO blocks LTP in the CA1 region of the hippocampus(Zhuo et al., 1993). However, HO-2 deficient mice have been reported to have normal hippocampal CA1 LTP(Poss et al., 1995). In contrast to CO, a role for NO in synaptic plasticity and learning has been previously reported(Muller, 1996; Balaban et al., 2014; Korshunova and Balaban, 2014). Thus, at this point it is an open question whether CO or NO evokes DA release in mammals. Downstream from CO or NO, RyRs have been shown to be required for hippocampal and cerebellum synaptic plasticity(Wang et al., 1996; Balschun et al., 1999; Lu and Hawkins, 2002; Kakizawa et al., 2012).

Our current results suggest that DA neurons release DA via two distinct mechanisms: canonical exocytosis and on-demand release. Canonical exocytosis is evoked by electrical activity of presynaptic DA neurons, requires Ca^2+^ influx, and may be involved in volume transmission. This mode of release can activate widespread targets over time, and is suited for regulating global brain functions. In contrast, on-demand release is evoked by activity of postsynaptic neurons, requires Ca^2+^ efflux via RyRs, and can regulate function of specific targets at precise times. DA neurons may differentially utilize these two modes of SV exocytosis in a context dependent manner. Understanding how DA neurons differentially utilize these modes of transmission will provide new insights into how a relatively small number of DA neurons can control numerous different brain functions.

## ACKNOWLEDGMENTS

This work was supported by grants from the Japanese Society for the Promotion of Science (JP17K07122) to K.U., a Grant-in-Aid for Scientific Research in Innovative Areas “Memory dynamism” (JP25115006) to M.S., and NIH (GM79465) to C.J.C. C.J.C. is an Investigator with the Howard Hughes Medical Institute. We thank A. Nose for *UAS-R-GECO1* transgenic flies, and T. Miyashita, M. Matsuno, T. Ueno and Y. Hirano for helpful discussions. The authors declare no competing financial interests.

## SUPPLEMENTAL INFROMATION

**Figure S1.**
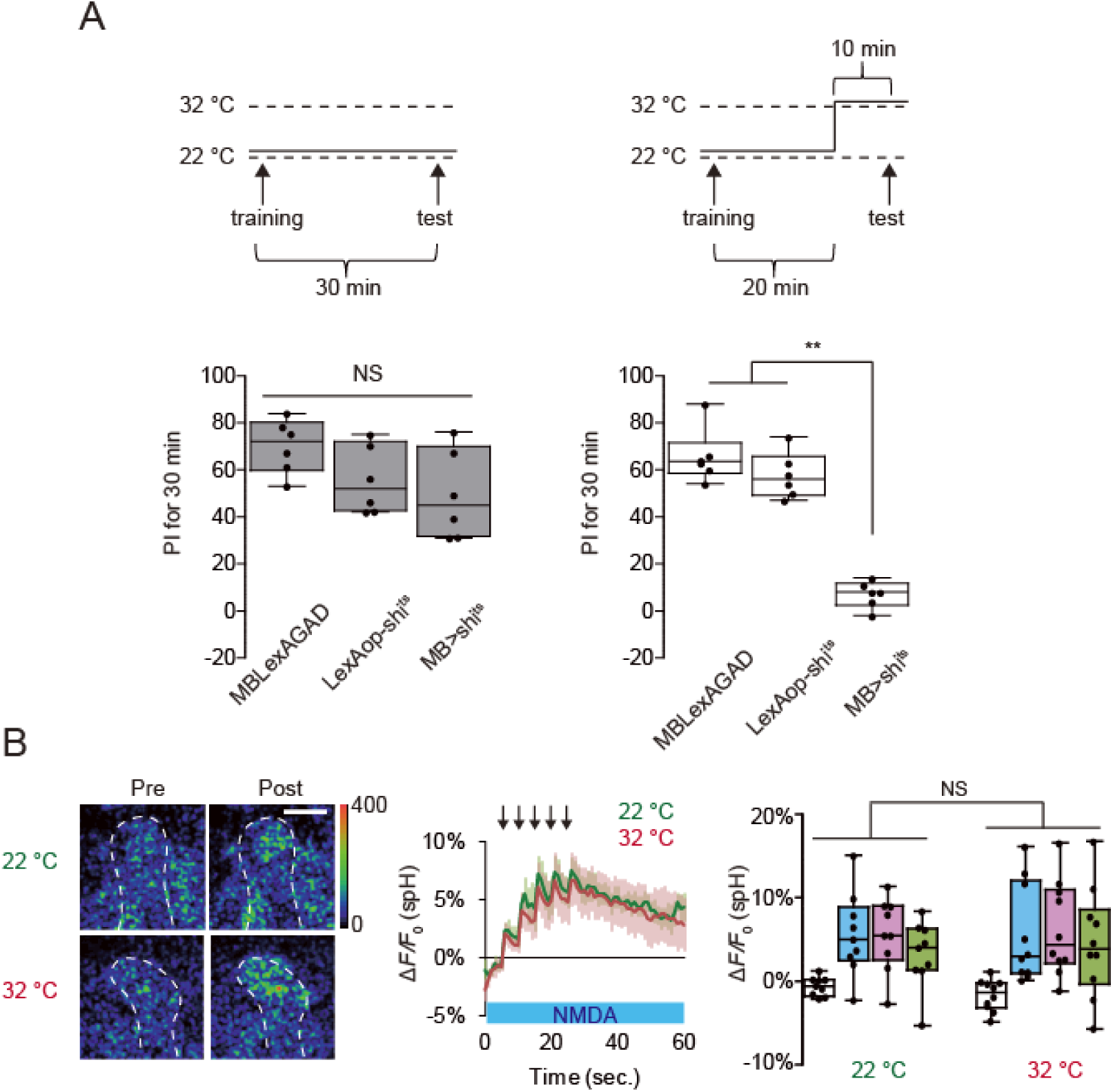
Coincident MB stimulation induces SV exocytosis from DA neurons. MB output is not required for DA release. **A**, Inhibiting MB output impairs recall of olfactory memory. One-way ANOVA and Bonferroni post hoc tests indicate significant impairment in memory recall in *MB > shi^ts^*flies at restrictive temperature (32°C) (*F_2,15_* = 71.05, *P<* 0.001) but not at permissive temperature (22°C) (*F_2,15_* = 2.854, *P = 2.854*). \*\**P* < 0.01 and NS *P* > 0.05. N = 6 for all data. **B**, Inhibiting MB output did not affect DA release. Two-way ANOVA identified no significant changes in spH fluorescence between restrictive (32 °C) and permissive temperatures (22 °C). N = 9 for 32 °C and N = 10 for 22°C.

**Figure S2.**
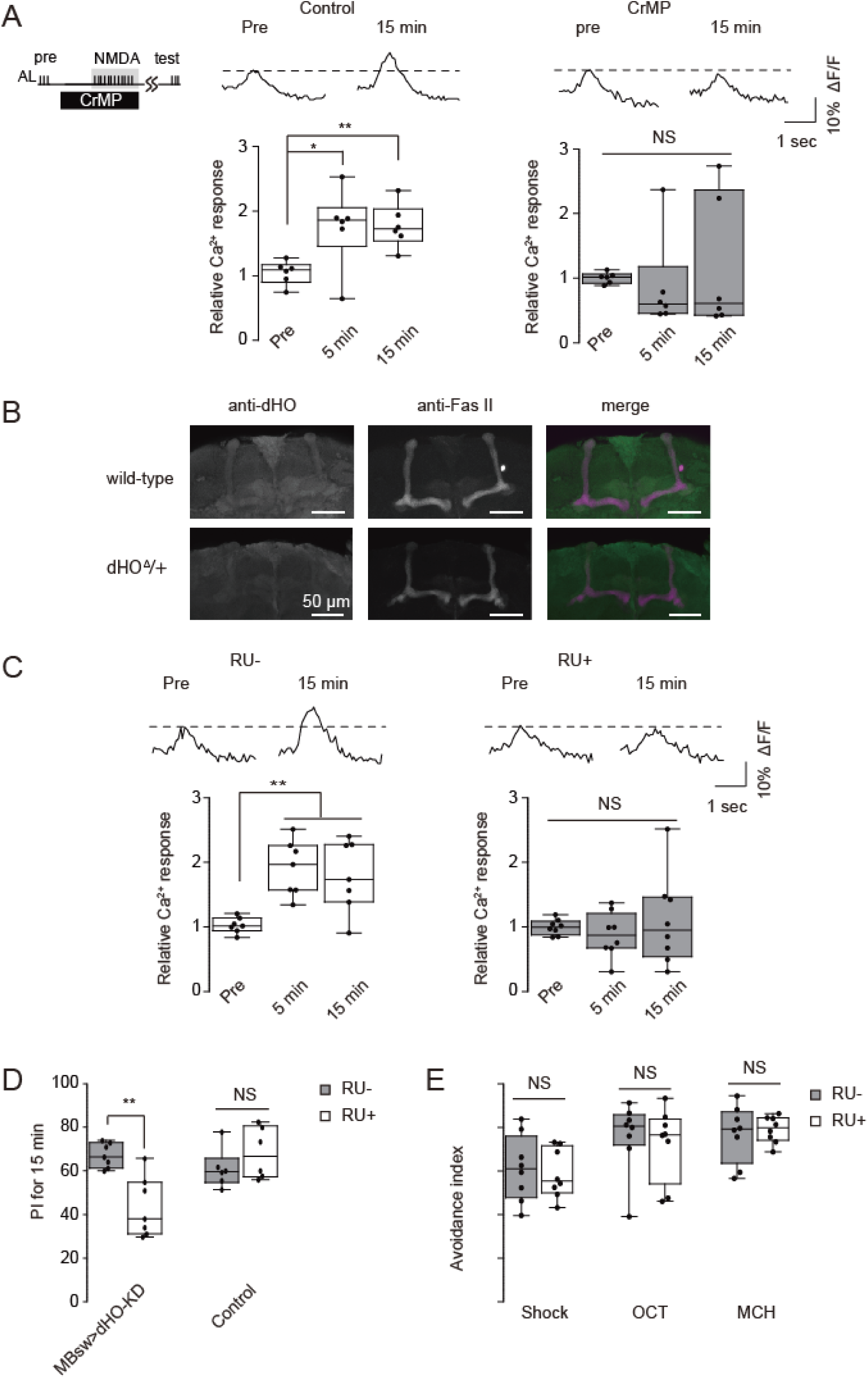
dHO in the MBs is required LTE and olfactory learning. **A**, Effects of HO inhibitor, CrMP, on LTE induced by AL+ NMDA stimulation. One-way ANOVA and Bonferroni post hoc tests indicate significant changes in AL-evoked Ca^2+^ responses in the MB after AL + NMDA stimulation in control conditions (*F_2,15_* = 5.836, *P* = 0.013, N = 6) and but not in 10 μM CrMP treated conditions (*F_2,18_* = 0.339, *P* = 0.717, N = 7). **B**, dHO antibody labels the MB lobes and midline cells in the *Drosophila* brain. Fas II antibody staining is included to identify subsets of the MB lobes. dHO signals are reduced in *dHO*^Δ^ hemizygotes (*dHO*^Δ^/+) demonstrating the specificity of the antibody. **C**, Knockdown of *dHO* in the MBs impairs LTE induced by AL + NMDA stimulation. One-way ANOVA and Bonferroni post hoc tests indicate significant LTE in the MB after AL + NMDA stimulation in control (RU-) conditions where *dHO* is not knocked down (Left panel, *F_2,18_*= 9.630, *P* = 0.001, N =7). LTE is not observed when dHO is knocked down (RU+) (Right panel, *F_2,21_*= 0.444, *P* = 0.647, N = 8). **D**, Knocking down *dHO* in the MBs impairs olfactory learning. \*\**P* < 0.01 determined by Student’s t-test. N = 7 for all data. **e**, Naïve responses to odors and electrical shock are not affected by knocking down *dHO* in the MBs. N = 8 for all experiments.

**Figure S3.**
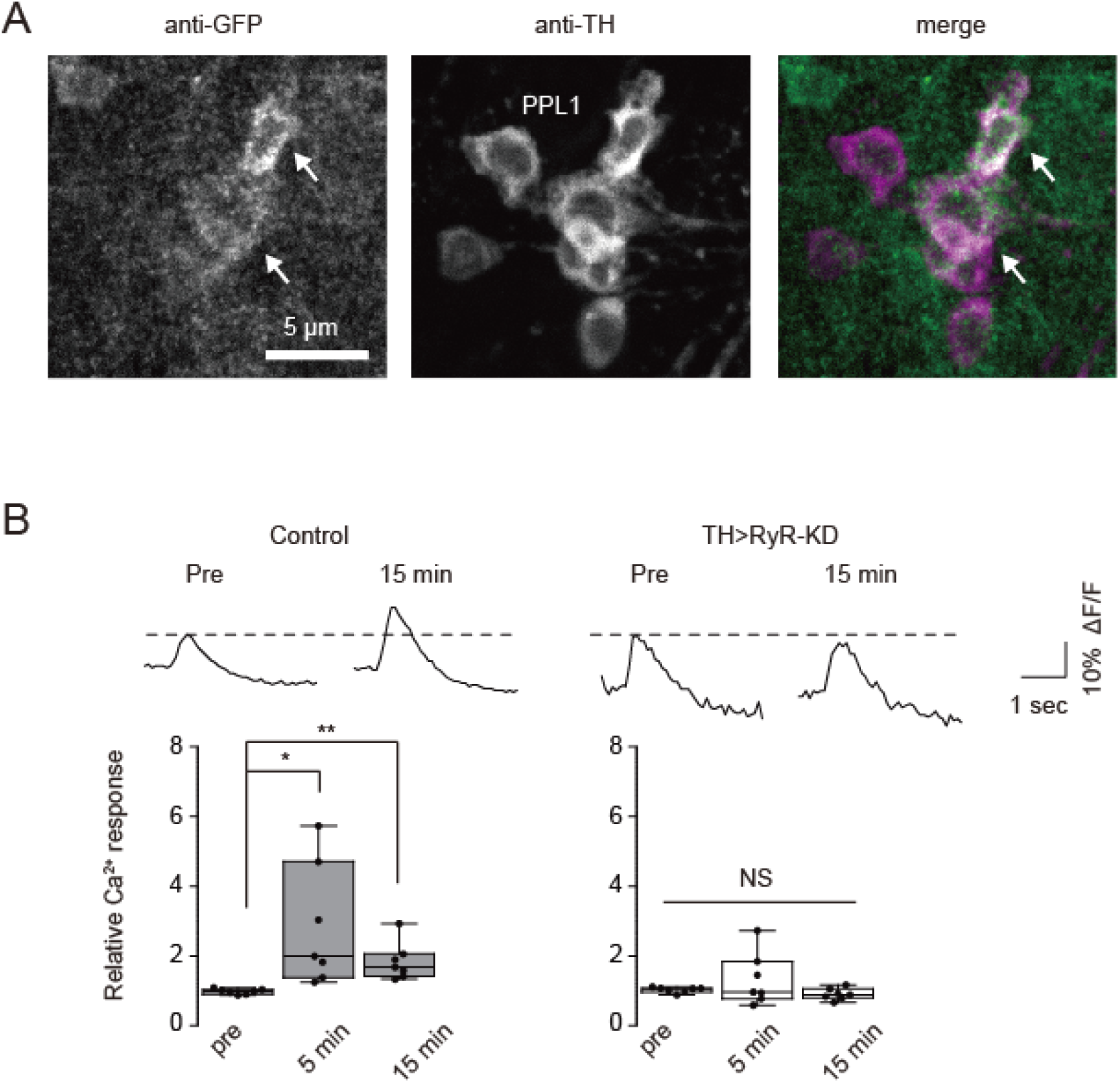
RyRs in TH-DA terminals are required for LTE. **A**, RyR localization was examined in *UAS-mCD8::GFP/Mi{Trojan-GAL4.0} RyR[MI08146-TG4.0]* flies in which expression of mCD8::GFP is driven by Trojan-GAL4 inserted in the endogenous *RyR* gene. GFP expression overlapped with expression of tyrosine hydroxylase (TH) in PPL1 DA neurons (arrows) that innervate the vertical lobes of the MBs. **B**, Knocking down RyRs in TH-DA neurons abolishes LTE induced by AL + NMDA stimulation. One-way ANOVA and Bonferroni post hoc tests indicate significant increases in AL-evoked Ca^2+^ responses after AL + NMDA stimulation in control brains (*F_2,18_*= 5.455, *P* = 0.014) but not in TH>RyR-KD brains (*F_2,18_*= 1.666, *P* = 0.217). N = 7 for all data.

**Figure S4.**
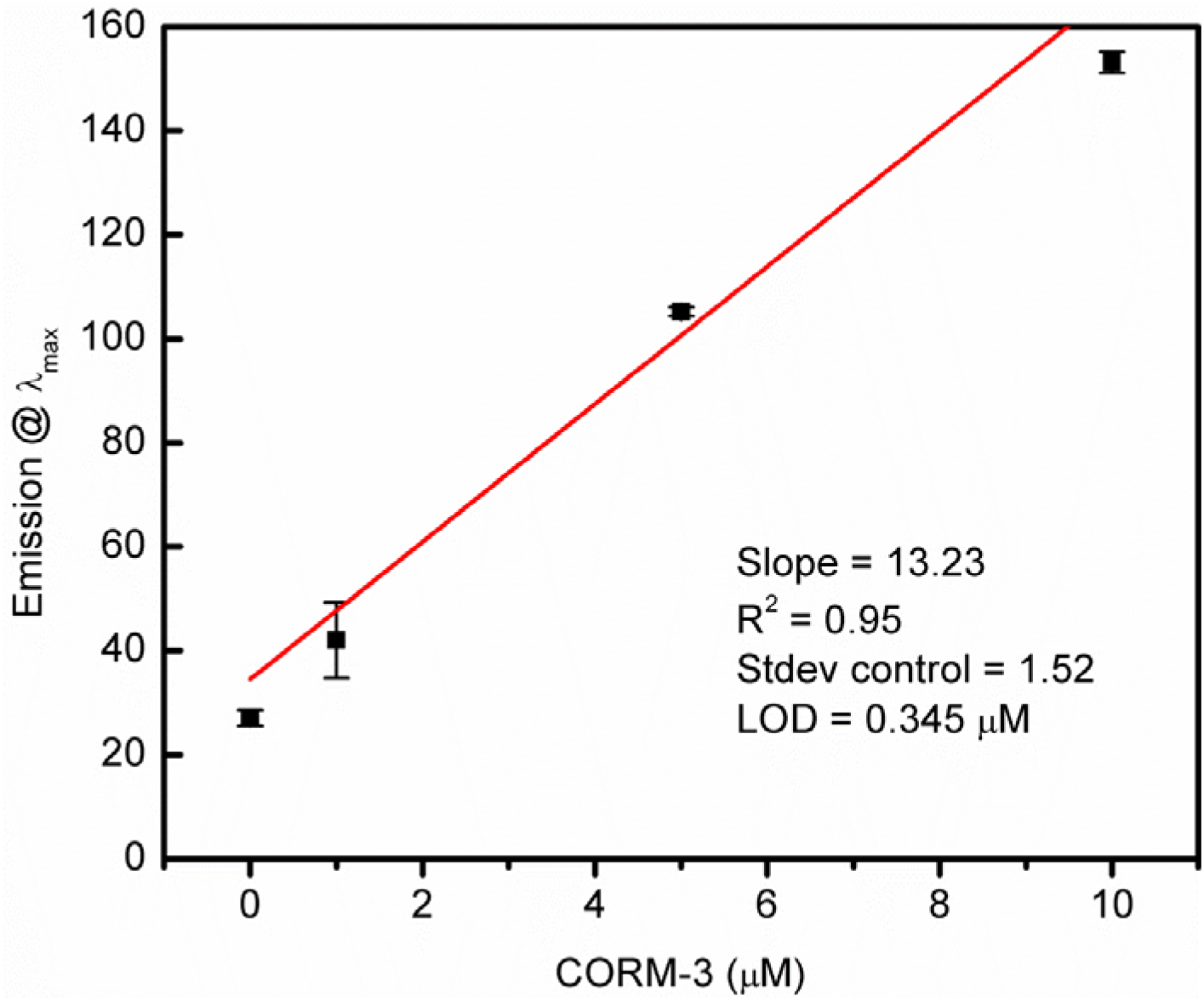
Limit of detection determination for COP-1. Limit of detection (LOD) of COP-1 at 90 min determined by fluorescence intensity at 508 nm as a function of CORM-3 concentration. While LOD of COP-1 is 0.345 μM at 90 min, a similar range of LOD’s were found at other timepoints (LOD = 0.660 μM at t = 30 min, 0.673 μM at t = 45 min, 0.556 μM at 60 min). Each concentration was run in triplicate and control was run five times. All data are shown as mean ± SD.

**Table S1.**
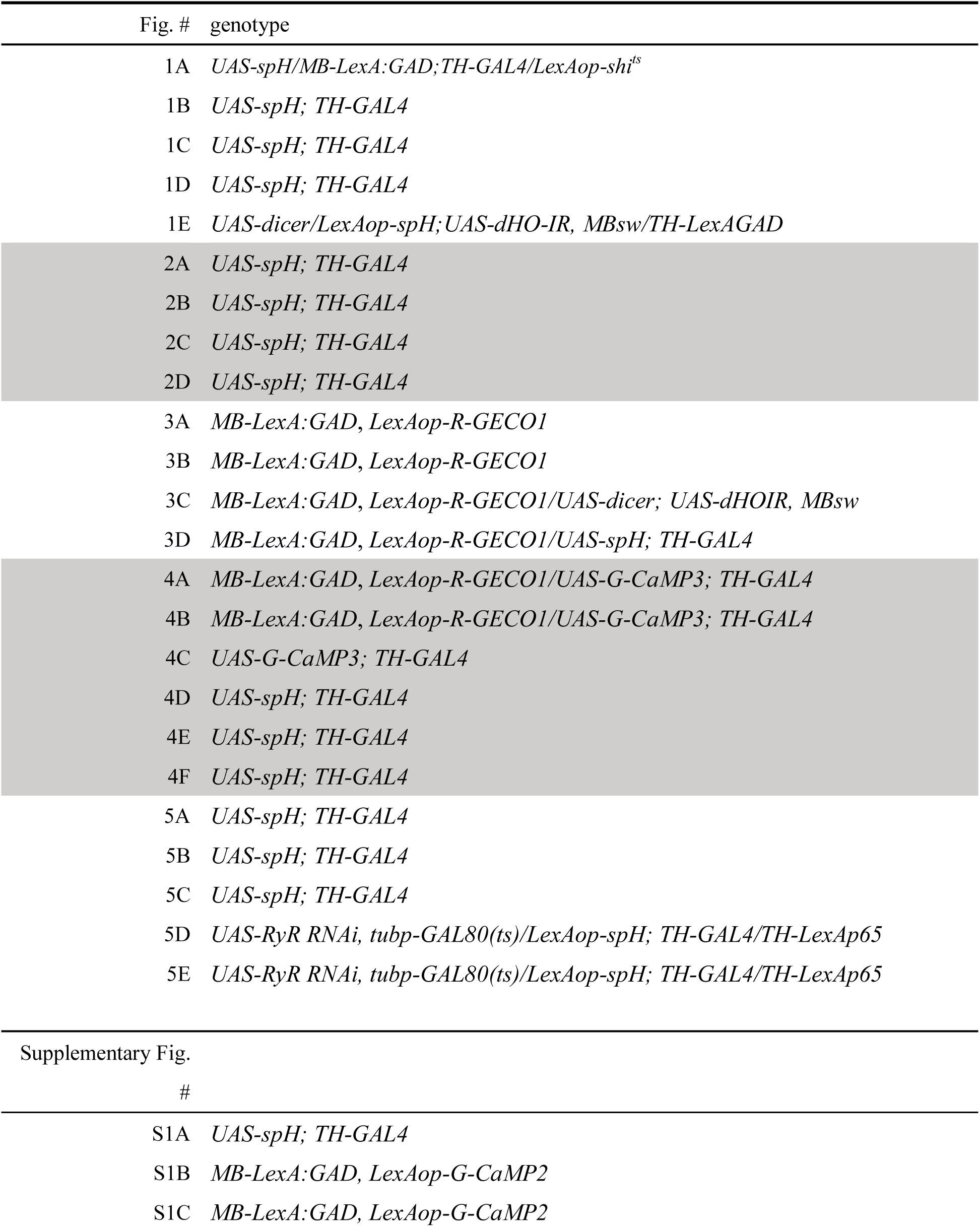
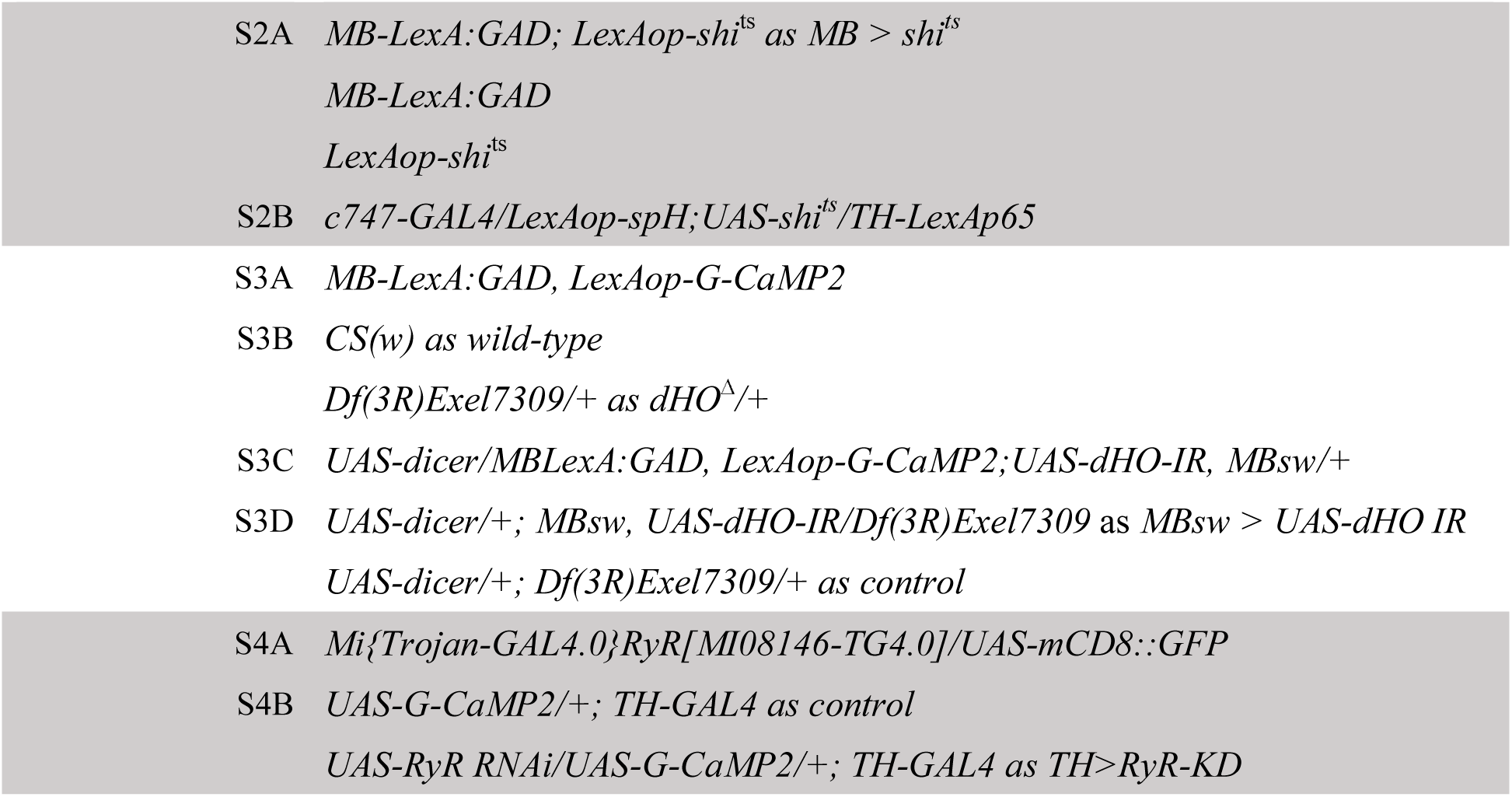
Genotypes used in each experiment

