## Supplemental Figures and Table for "Carbon monoxide, a retrograde messenger generated in post-synaptic mushroom body neurons evokes non-canonical dopamine release"

**This supplemental Infromation includes:**

Figures S1 to S4

Table S1

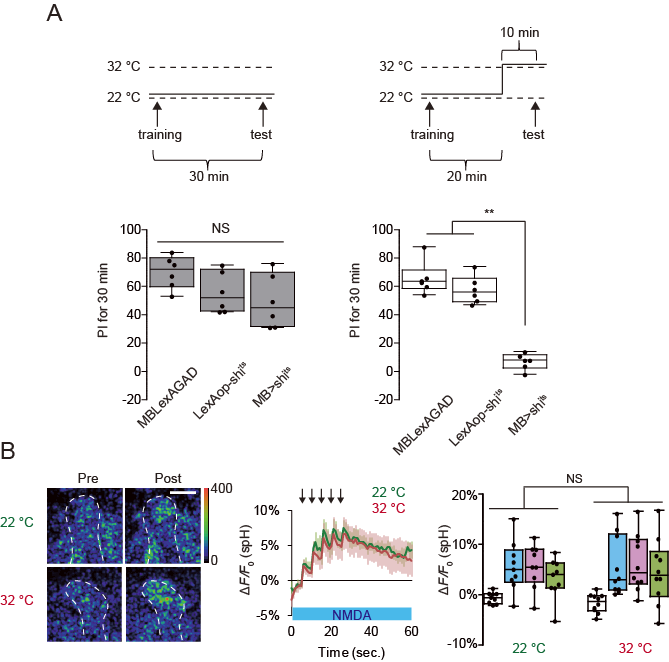

**Figure S1 Coincident MB stimulation induces SV exocytosis from DA neurons. MB output is not required for DA release.**

**A,** Inhibiting MB output impairs recall of olfactory memory. One-way ANOVA and Bonferroni post hoc tests indicate significant impairment in memory recall in *MB > shi^ts^* flies at restrictive temperature (32°C) (*F_2,15_* = 71.05, *P<* 0.001) but not at permissive temperature (22°C) (*F_2,15_* = 2.854, *P = 2.854*). ***P* < 0.01 and NS *P* > 0.05. N = 6 for all data. **B**, Inhibiting MB output did not affect DA release. Two-way ANOVA identified no significant changes in spH fluorescence between restrictive (32 °C) and permissive temperatures (22 °C). N = 9 for 32 °C and N = 10 for 22°C.

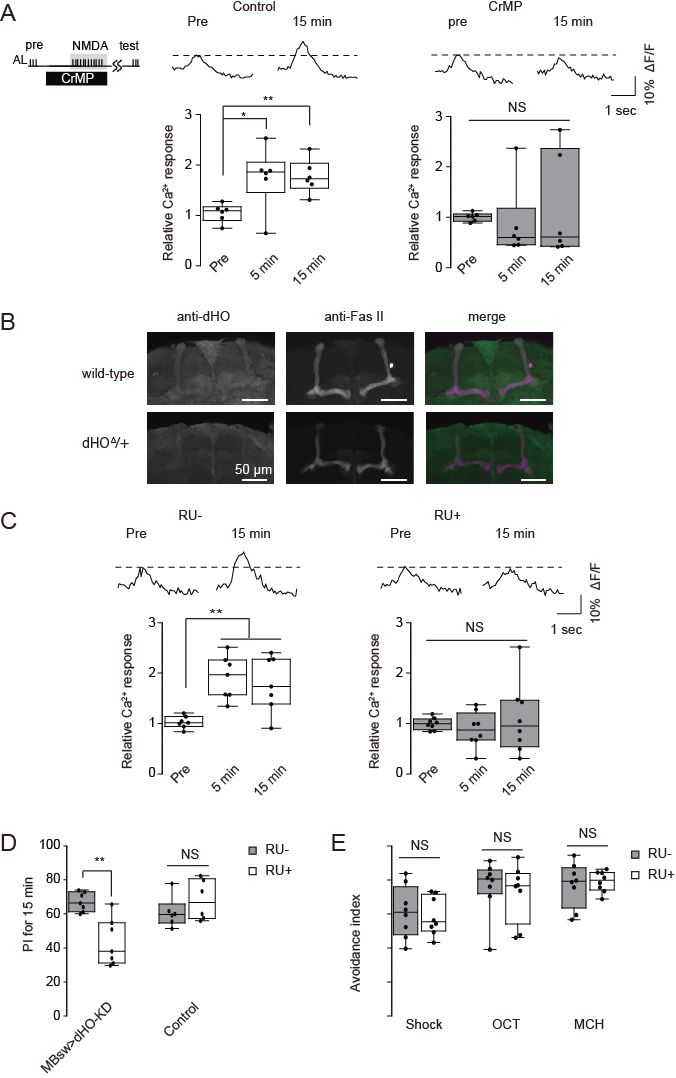

**Figure S2 dHO in the MBs is required LTE and olfactory learning.**

**A**, Effects of HO inhibitor, CrMP, on LTE induced by AL+ NMDA stimulation. One-way ANOVA and Bonferroni post hoc tests indicate significant changes in AL-evoked Ca^2+^ responses in the MB after AL + NMDA stimulation in control conditions (*F_2,15_* = 5.836, *P* = 0.013, N = 6) and but not in 10 μM CrMP treated conditions (*F_2,18_* = 0.339, *P* = 0.717, N = 7). **B**, dHO antibody labels the MB lobes and midline cells in the *Drosophila* brain. Fas II antibody staining is included to identify subsets of the MB lobes. dHO signals are reduced in *dHO^Δ^* hemizygotes (*dHO^Δ^*/+) demonstrating the specificity of the antibody. **C**, Knockdown of *dHO* in the MBs impairs LTE induced by AL + NMDA stimulation. One-way ANOVA and Bonferroni post hoc tests indicate significant LTE in the MB after AL + NMDA stimulation in control (RU-) conditions where *dHO* is not knocked down (Left panel, *F_2,18_*= 9.630, *P* = 0.001, N =7). LTE is not observed when dHO is knocked down (RU+) (Right panel, *F_2,21_*= 0.444, *P* = 0.647, N = 8). **D**, Knocking down *dHO* in the MBs impairs olfactory learning. ***P* < 0.01 determined by Student's t-test. N = 7 for all data. **e**, Naïve responses to odors and electrical shock are not affected by knocking down *dHO* in the MBs. N = 8 for all experiments.

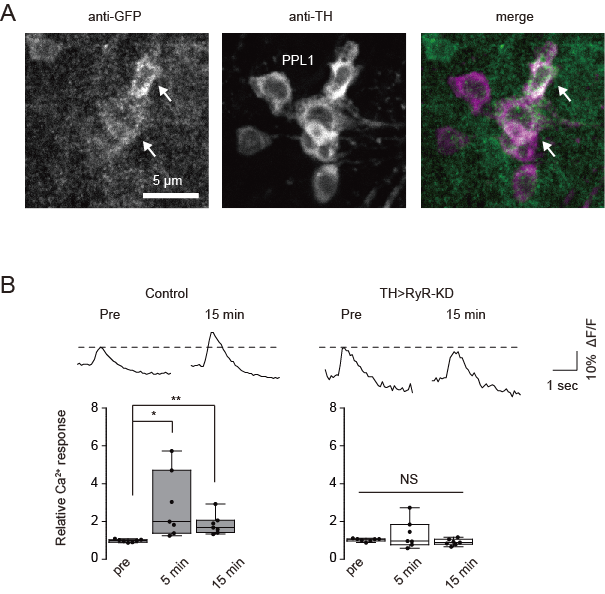

**Figure S3 RyRs in TH-DA terminals are required for LTE.**

**A**, RyR localization was examined in *UAS-mCD8::GFP/Mi{Trojan-GAL4.0} RyR[MI08146-TG4.0]* flies in which expression of mCD8::GFP is driven by Trojan-GAL4 inserted in the endogenous *RyR* gene. GFP expression overlapped with expression of tyrosine hydroxylase (TH) in PPL1 DA neurons (arrows) that innervate the vertical lobes of the MBs. **B**, Knocking down RyRs in TH-DA neurons abolishes LTE induced by AL + NMDA stimulation. One-way ANOVA and Bonferroni post hoc tests indicate significant increases in AL-evoked Ca^2+^ responses after AL + NMDA stimulation in control brains (*F_2,18_*= 5.455, *P* = 0.014) but not in TH>RyR-KD brains (*F_2,18_*= 1.666, *P* = 0.217). N = 7 for all data.

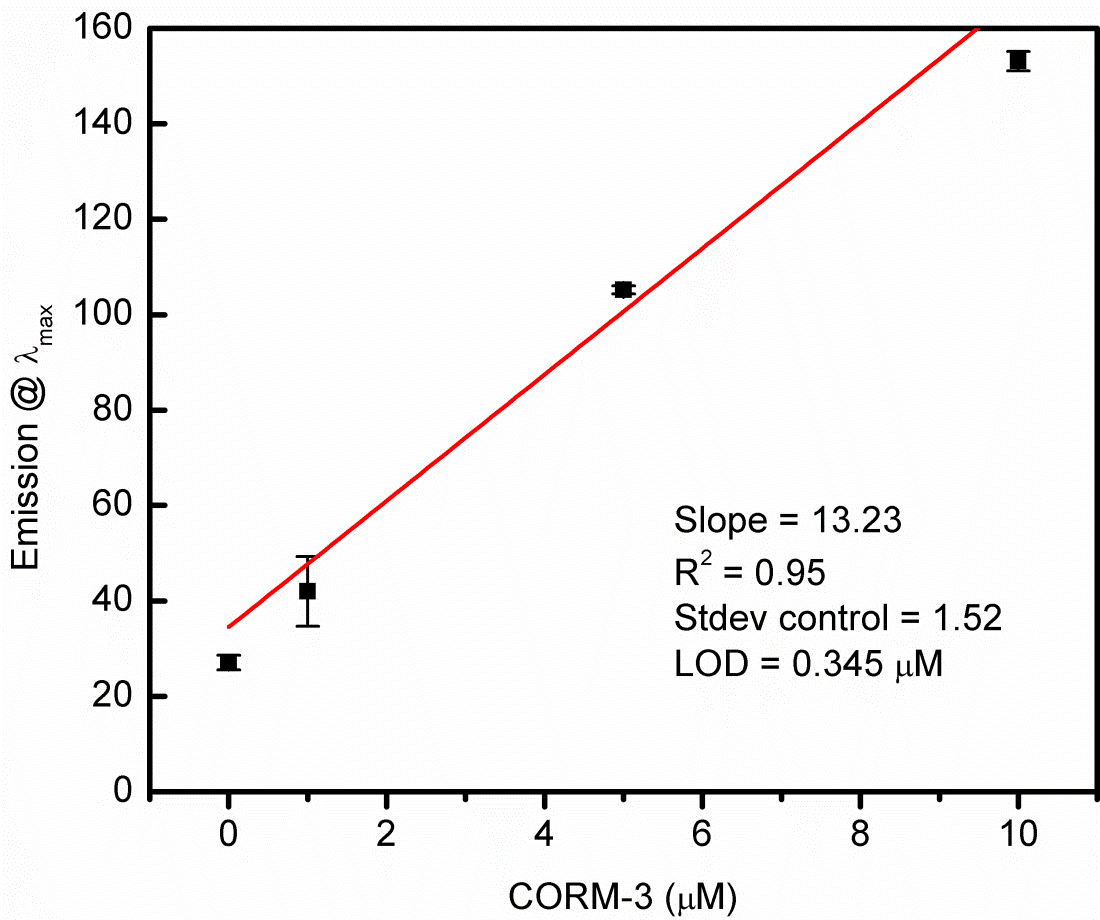

**Figure S4 Limit of detection determination for COP-1.**

Limit of detection (LOD) of COP-1 at 90 min determined by fluorescence intensity at 508 nm as a function of CORM-3 concentration. While LOD of COP-1 is 0.345 μM at 90 min, a similar range of LOD’s were found at other timepoints (LOD = 0.660 μM at t = 30 min, 0.673 μM at t = 45 min, 0.556 μM at 60 min). Each concentration was run in triplicate and control was run five times. All data are shown as mean ± SD.

| Table S1 Genotypes used in each experiment | |
| --- | --- |
| Fig. # | genotype |
| 1A | *UAS-spH/MB-LexA:GAD;TH-GAL4/LexAop-shi^ts^* |
| 1B | *UAS-spH; TH-GAL4* |
| 1C | *UAS-spH; TH-GAL4* |
| 1D | *UAS-spH; TH-GAL4* |
| 1E | *UAS-dicer/LexAop-spH;UAS-dHO-IR, MBsw/TH-LexAGAD* |
| 2A | *UAS-spH; TH-GAL4* |
| 2B | *UAS-spH; TH-GAL4* |
| 2C | *UAS-spH; TH-GAL4* |
| 2D | *UAS-spH; TH-GAL4* |
| 3A | *MB-LexA:GAD*, *LexAop-R-GECO1* |
| 3B | *MB-LexA:GAD*, *LexAop-R-GECO1* |
| 3C | *MB-LexA:GAD*, *LexAop-R-GECO1/UAS-dicer; UAS-dHOIR, MBsw* |
| 3D | *MB-LexA:GAD*, *LexAop-R-GECO1/UAS-spH; TH-GAL4* |
| 4A | *MB-LexA:GAD*, *LexAop-R-GECO1/UAS-G-CaMP3; TH-GAL4* |
| 4B | *MB-LexA:GAD*, *LexAop-R-GECO1/UAS-G-CaMP3; TH-GAL4* |
| 4C | *UAS-G-CaMP3; TH-GAL4* |
| 4D | *UAS-spH; TH-GAL4* |
| 4E | *UAS-spH; TH-GAL4* |
| 4F | *UAS-spH; TH-GAL4* |
| 5A | *UAS-spH; TH-GAL4* |
| 5B | *UAS-spH; TH-GAL4* |
| 5C | *UAS-spH; TH-GAL4* |
| 5D | *UAS-RyR RNAi, tubp-GAL80(ts)/LexAop-spH; TH-GAL4/TH-LexAp65* |
| 5E | *UAS-RyR RNAi, tubp-GAL80(ts)/LexAop-spH; TH-GAL4/TH-LexAp65* |
| Supplementary Fig. # |  |
| S1A | *UAS-spH; TH-GAL4* |
| S1B | *MB-LexA:GAD, LexAop-G-CaMP2* |
| S1C | *MB-LexA:GAD, LexAop-G-CaMP2* |
| S2A | *MB-LexA:GAD; LexAop-shi*^ts^ *as MB > shi^ts^* |
|  | *MB-LexA:GAD* |
|  | *LexAop-shi*^ts^ |
| S2B | *c747-GAL4/LexAop-spH;UAS-shi^ts^/TH-LexAp65* |
| S3A | *MB-LexA:GAD, LexAop-G-CaMP2* |
| S3B | *CS(w) as wild-type* |
|  | *Df(3R)Exel7309/+ as dHO*^∆^*/+* |
| S3C | *UAS-dicer/MBLexA:GAD, LexAop-G-CaMP2;UAS-dHO-IR, MBsw/+* |
| S3D | *UAS-dicer/+; MBsw, UAS-dHO-IR/Df(3R)Exel7309* as *MBsw > UAS-dHO IR* |
|  | *UAS-dicer/+; Df(3R)Exel7309/+* *as control* |
| S4A | *Mi{Trojan-GAL4.0}RyR[MI08146-TG4.0]/UAS-mCD8::GFP* |
| S4B | *UAS-G-CaMP2/+; TH-GAL4 as control* |
|  | *UAS-RyR RNAi/UAS-G-CaMP2/+; TH-GAL4 as TH>RyR-KD* |
